# Inter-population variation in the Atlantic salmon microbiome reflects environmental and genetic diversity

**DOI:** 10.1101/283754

**Authors:** Tamsyn M. Uren Webster, Sofia Consuegra, Matthew Hitchings, Carlos Garcia de Leaniz

## Abstract

Microbial communities have a crucial influence on host phenotype, and are of broad interest to ecological and evolutionary research. Yet, the extent of variation that occurs in the microbiome within and between populations is unclear. We characterised the skin and gut microbiome of seven populations of juvenile Atlantic salmon (*Salmo salar*) inhabiting a diverse range of environments, including hatchery-reared and wild populations. We found shared skin OTUs across all populations and core gut microbiota for all wild fish, but the diversity and structure of both skin and gut microbial communities were distinct between populations. There was a marked difference between the gut microbiome of wild and captive fish. Hatchery-reared fish had lower intestinal microbial diversity, lacked core microbiota found in wild fish, and showed altered community structure and function. Captive fish skin and gut microbiomes were also less variable within populations, reflecting more uniform artificial rearing conditions. Surrounding water influenced the microbiome of the gut and, especially, the skin, but could not explain the degree of variation observed between populations. For both the gut and the skin, we found that there was greater difference in microbial community structure between more genetically distinct fish populations, and also that population genetic diversity was positively correlated with microbiome diversity. However, dietary differences are likely to be the major factor contributing to the large differences found in the gut microbiome between wild and captive fish. Our results highlight the scope of inter-population variation in the microbiome, and offer insights into the contributing deterministic factors.

## Introduction

There is increasing recognition that host-associated microbial communities have a fundamental effect on organism phenotype; aiding digestion and nutrient acquisition, influencing metabolism, energy storage, growth and behaviour, and playing a critical role in immune system maturation and pathogen defence (Hird 2017; Hooper et al. 2012; Koskella et al. 2017). Microbial communities are very dynamic and have an extensive capacity to respond to local selective pressures via phenotypic plasticity, rapid mutation rates, short generation times and high intra-community gene flow (Walter & Ley 2011). Such metagenomic plasticity has been proposed to also specifically enhance phenotypic plasticity in the host, for example through improving thermoregulation capacity, enabling digestion of novel food sources or increased resistance to local pathogens, and may contribute to host acclimation or even population-level adaptation to environmental change (Alberdi et al. 2016; Walter & Ley 2011). The microbiome is therefore of great interest to many aspects of ecological and evolutionary research, including questions related to potential drivers of local adaptation and domestication, and in assessing the likely impacts of environmental stressors. However, the precise mechanistic drivers of microbiome community dynamics have yet to be established (Alberdi et al. 2016; Hird 2017; Koskella et al. 2017), especially for non-mammalian species including fish, which also experience distinct ecological interactions within the aquatic environment (Llewellyn et al. 2014).

The structure and diversity of the vertebrate microbiome is determined by complex and dynamic interactions between the host, the local environment, and other microbiota (Walter & Ley 2011). Deterministic factors, including host-specific and environmental filters, are thought to be the dominant forces shaping teleost microbial communities, with stochastic processes playing a minor role (Roeselers et al. 2011; Schmidt et al. 2015; Sullam et al. 2015). Initial microbial colonisation of the teleost intestine after hatching is thought to be seeded by microbes in the surrounding water (Giatsis et al. 2015; Ingerslev et al. 2014). Upon first feeding, diet becomes a dominant factor in shaping further proliferation and differentiation of the gut microbiome, and dietary changes can alter the diversity and structure of gut microbial communities throughout life (Gajardo et al. 2016; Giatsis et al. 2015; Smith et al. 2015). Water temperature (Zarkasi et al. 2014), pH (Sylvain et al. 2016), salinity (Llewellyn et al. 2016; Schmidt et al. 2015) and habitat type (Smith et al. 2015; Sullam et al. 2015), as well as developmental stage (Llewellyn et al. 2016), sex (Li et al. 2016), the immune system (Bolnick et al. 2014; Stagaman et al. 2017) and genetic background (Smith et al. 2015; Sullam et al. 2015) have been shown to affect the diversity and/or structure of the teleost intestinal microbiome. Inter-host dispersal of microbiota has recently also been shown to make a significant contribution to intestinal microbiome variation, at least in laboratory-reared zebrafish (Burns et al. 2017). Less research has focused on microbial communities associated with other mucosal surfaces, but pH (Sylvain et al. 2016), stress (Boutin et al. 2013) and salinity (Lokesh & Kiron 2016) have been shown to influence fish skin microbial communities.

Despite these known effects of environmental and host-specific factors, very little is known about the degree of variation in microbial community diversity and structure that occurs within and between fish populations. Variation in the microbiome is likely to have a fundamental role in host ecology and evolution (Hird 2017; Suzuki 2017). Knowledge about the degree and nature of microbiome variation, and the mechanistic drivers behind it, is crucial for understanding how host-associated microbial communities may influence host phenotype and contribute to evolutionary processes (Suzuki 2017). To date, most vertebrate microbiome research has focused on captive-reared individuals or laboratory animals, which may not be representative of the full extent of natural microbiome variation, therefore microbial profiling of natural populations is a priority (Colston & Jackson 2016; Hird 2017; Rosshart et al. 2017). Furthermore, the vast majority of studies have focused on the intestinal microbiome, despite the fact that other mucosal surfaces, such as the skin, are also likely to have a critical influence of host health and are likely to be under different selective pressures (Llewellyn et al. 2014).

We therefore aimed to examine the degree and nature of variation in the gut and skin microbiome across seven diverse populations of juvenile Atlantic salmon (*Salmo salar*), including wild and hatchery-reared fish. We also examined the potential role of host genetic background and environmental factors in contributing to this variation. Atlantic salmon is an excellent model for investigating variation in the microbiome because it is a locally adapted species which experiences a high degree of environmental variation and shows substantial genetic differentiation among natural populations (Garcia de Leaniz et al. 2007). Atlantic salmon are also one of the more extensively domesticated finfish species, and are subjected to distinct evolutionary pressures in captivity due to artificial diets, water treatment and high stocking densities (Stringwell et al. 2014; Teletchea & Fontaine 2014). We hypothesised that wild and hatchery-reared fish would have distinct gut and skin microbial communities and, specifically, that captive salmon would have a less diverse microbiome. We also predicted that fish from each different population would have distinct microbiota, reflecting genetic and environmental diversity.

## Materials and methods

### Sample collection

Atlantic salmon fry (0+; all approximately 8-9 months post hatch) were sampled from four wild populations (rivers Spey, Tweed, Towy, and Frome) and three hatcheries in Scotland, Wales, England and France between October and December 2015 (Fig 1). We specifically selected geographically and geologically distinct sites to include environmental and genetic variation in this study. Twelve fish were sampled per population; wild fish were captured by electrofishing, while hatchery fish were captured by hand-netting. Fish were euthanised via an overdose of anaesthesia (Phenoxyethanol (0.5 mg/L), followed by destruction of the brain according to UK Home Office regulations. Skin mucus was sampled from each fish by swabbing the left-hand side of the body along the entire length of the lateral line 5x in both directions (Epicentre Catch-All™ Sample Collection Swabs, Cambio, Cambridge, UK). Gut samples were obtained by sterile dissection of the whole intestine to include both intestinal contents and epithelial associated microbial communities. Fifty mL water samples were also taken from each site. All samples were collected into sterile tubes, transported on dry ice and stored at −80 °C until DNA extraction.

**Figure 1.**
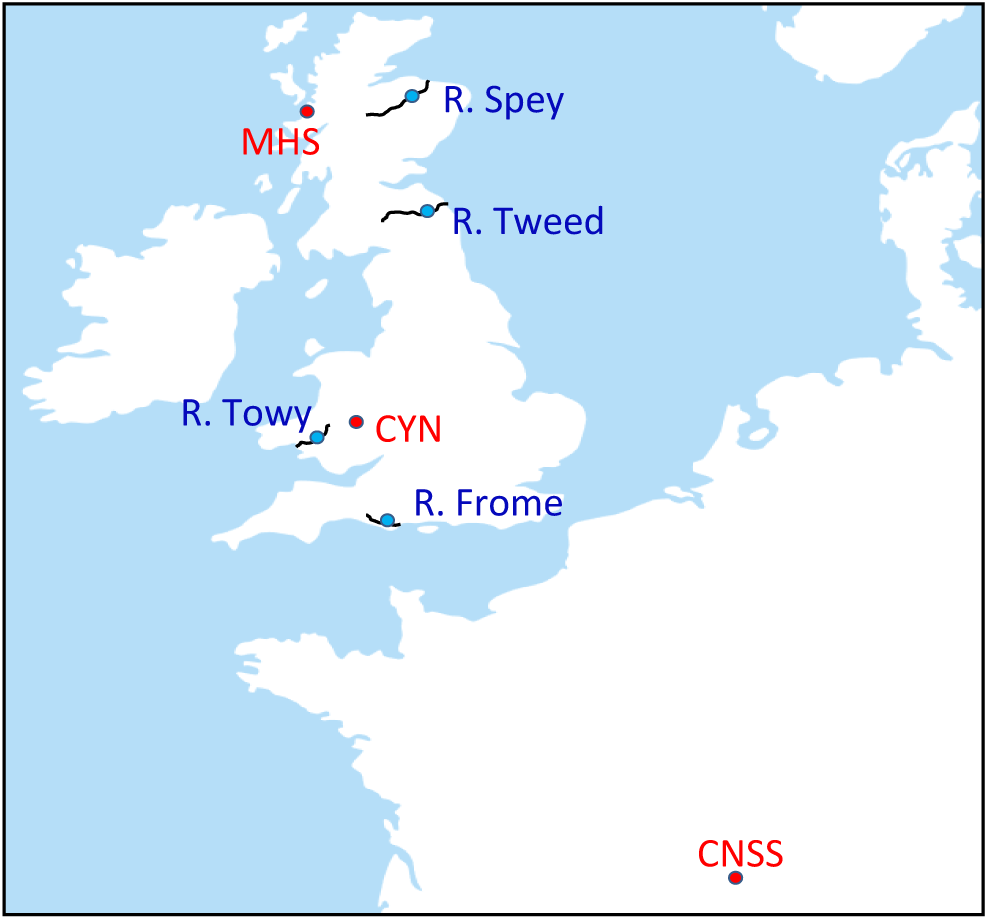
Location of study Atlantic salmon populations sampled as 0+ juveniles in four rivers (blue circles) and in three hatcheries (red circles).

Fork length and wet weight were recorded *in situ* and used to calculate Fulton’s factor as a measure of body condition. As sex is difficult to detect visually in young salmonids, we used a genetic marker for sex identification (Yano et al. 2013) to account for potential effects on microbial community structure.

### 16S rRNA amplicon sequencing

Briefly, DNA extraction from all gut, skin swab and water samples was performed using MoBio PowerSoil® DNA Isolation Kit (Cambio, Cambridge, UK). 16S library preparation was performed according to the Illumina Metagenomic Sequencing Library Preparation protocol (Illumina 2013), amplifying the V4 hypervariable region using primers selected as the best candidates for bacterial and archaeal representation (Klindworth et al. 2013); 519F (5’-CAGCMGCCGCGGTAA) and 785R (5’-TACNVGGGTATCTAATCC), using 12.5 ng total genomic input DNA and Nextera XT indexing. Purified PCR products were pooled in equal concentrations, before sequencing using an Illumina MiSeq (300 bp PE reads). Full experimental details are given in the supporting information. Two extraction blanks were prepared and sequenced together with the samples in order to assess the degree of background microbial contamination.

### Population genetics analysis

Each individual (12 fish/site) was genotyped at 12 neutral microsatellites DNA loci described by Ellis et al.(2011), together with two markers tightly linked to expressed MHC genes, embedded in the 3’ untranslated regions of MHCI and MHCII (*Sasa-UBA* and *Sasa-DAA*, respectively (Grimholt et al. 2003)), further details on reaction conditions are given in the supporting information. Fragment sizes were analysed using an ABI3130xl Genetic Analyser and estimated using GENEMAPPER 4.0 software (Applied Biosystems, Sussex, UK) using a GS LIZ 500(−150) size standard. Pairwise Nei’s genetic distances were calculated between populations and between all individuals, and values were additionally analysed using principal component analysis, using GenAlEx 6.5 (Peakall & Smouse 2012). Pairwise *F*_ST_ values between populations were calculated using Arlequin v.3.5.2.2 (Excoffier & Lischer 2010). Individual and population level observed heterozygosity was calculated using Cernicalin (Aparicio et al. 2006) and allelic richness calculated using FSTAT v2.9.3 (Goudet 1995).

### 16S rRNA bioinformatics analysis

Microbial community analysis was performed using mother v1.37 (Kozich et al. 2013), Qiiime v1.9 (Caporaso et al. 2010) and R v3.3.2 (R CoreTeam 2014), with full details given in the supporting information. Briefly raw reads were quality filtered using Trimmomatic (Bolger et al. 2014), merged, filtered and aligned to the Silva seed reference database (version 123) (Quast et al. 2013). Potential chimeras were removed using UCHIME (Edgar et al. 2011) before taxonomic classification using the Silva reference taxonomy and removal of mitochondrial, eurkaryote and chloroplast sequences.

Phylogenetic trees were constructed using Clearcut (Evans et al. 2006), then weighted Unifrac (Lozupone et al. 2011) distances between samples were calculated and used for analysis of microbial community structure within mothur. Non-metric multidimensional scaling analysis was used for structural visualisation. Statistical analysis of Unifrac distances was performed using Adonis in the Vegan package in R (Oksanen et al. 2017) to assess the effects of origin (wild/hatchery), population, fork length, condition factor, sex, individual heterozygosity and individual MHC heterozygosity on community structural variance, using the strata function to specify a nested model of population within group origin. Additionally, HOMOVA, within mothur, was used to specifically quantify the degree of intra-population variation in community structure. A Mantel test was employed to test for correlation between individual level genetic distances and weighted Unifrac distances for both the gut and the skin.

Analysis of microbial community alpha diversity and composition were performed at the operational taxonomic unit (OTU) level, based on 97% sequence similarity. In order to maximise sample inclusion, whilst ensuring high Good’s coverage (≥ 94%) for all included samples, reads were subsampled to a depth of 4012/sample, retaining 76 gut and 81 skin samples (minimum of 10/population), together with a water sample from each site. We calculated two measures of alpha diversity (Chao1 richness and Shannon diversity) using mothur. Variation in alpha diversity was analysed by linear mixed modelling with the *lme4* package in R using origin (hatchery/wild), length, condition factor, sex, individual heterozygosity and individual MHC heterozygosity as fixed factors, and population as a random factor to account for spatial autocorrelation. Water microbial diversity at each site was included as an *offset* to statistically control for the effects of the surrounding water on fish microbial diversity.

OTUs that were present in at least 80% of all individuals were identified using the compute_core_microbiome function in Qiime. Following filtering of singleton OTUs, a Kruskal-Wallace test, incorporating FDR correction, was implemented using the group_significance function in Qiime to identify differentially abundant OTUs between wild and hatchery-origin fish. In addition to these OTU-level analyses, further community composition analysis was performed at the Phylum level using the summarize_taxa function in Qiime, followed by cluster analysis of all gut, skin and water samples using the Bray-Curtis similarity index. Additionally, functional analysis of community structure based on ‘metabolism’ was performed using OTU taxon to phenotype mapping with METAGENassist (Arndt et al. 2012) and visualised using Euclidean distance clustering.

## Results

### Phenotypic and genetic differentiation among study populations

As expected, hatchery-reared fish were on average larger (*t*_45.288_ = 5.40, *P*< 0.001), heavier (*t*_37.044_ = 5.09, *P*< 0.001) and had a higher condition factor, K (*t*_81.791_ = 4.30, *P*< 0.001) than wild fish of the same age, although there were also significant differences among populations (length, *F*_6,78_ = 217.53, *P*< 0.001; mass, *F*_6,78_ = 183.93, *P*< 0.001; K, *F*_6,78_ = 8.73, *P*< 0.001). Observed heterozygosity calculated using MHC-linked, but not neutral, loci was significantly higher in wild than in hatchery populations (MHC-He: t= 3.25, *P*= 0.02; neutral-He: t= 0.57, *P*= 0.59), but there were no significant differences in allelic richness ((MHC-AR: t= 2.43, *P*= 0.06; neutral-AR: t= 2.16, *P*= 0.08; Table S1). Pairwise *F*_ST_ values were significant for all comparisons (*P* < 0.01), except for the two Scottish rivers (Tweed and Spey; Table S2), although there was considerable variation in *F*_ST_ and genetic distances between different populations. Genetic differentiation of populations is also highlighted by the results from PCoA analysis of genetic distance (Figure S1).

### Structural analysis of microbial communities

After quality filtering, database alignment, and removal of potential chimeras, mitochondrial, chloroplast and eukaryotic sequences, a total of 7,254,876 good quality sequences were retained for analysis. Fewer than 30 bacterial sequences were identified from each of the extraction blanks confirming little impact of background contamination.

NMDS cluster analysis of weighted Unifrac distances for all individuals, incorporating sequence phylogeny and relative abundance, for both the skin and the gut is shown in Figure 2. For the gut, there was a significant effect of both origin (hatchery/wild) and population on community structure, but not sex, fork length, condition factor or individual heterozygosity or MHC heterozygosity (origin; F_1_=4.15, *P=*0.002, popn; F_6_=2.57, *P=*0.001). For the skin, there was also a significant effect of origin and population, as well as fork length, on community structure (origin; F_1_=4.28, *P*=0.001, popn; F_6_=6.77, *P=0*.001, length; F_1_=2.72, *P*=0.01). There was a significant correlation between individual weighted Unifrac distances and genetic distances (calculated using all 14 markers) for both the gut (M=0.20, *P*=0.002) and the skin (M=0.35, *P*<0.001). Analysis of homogeneity of structural variance between populations using HOMOVA revealed markedly higher inter-individual variance in all wild populations compared to the hatchery populations (gut: t_5_=6.55, *P=*0.001, skin: t_5_=4.60, *P=*0.006), while the skin microbiome was generally more homogenous between individuals than the gut microbiome.

**Figure 2.**
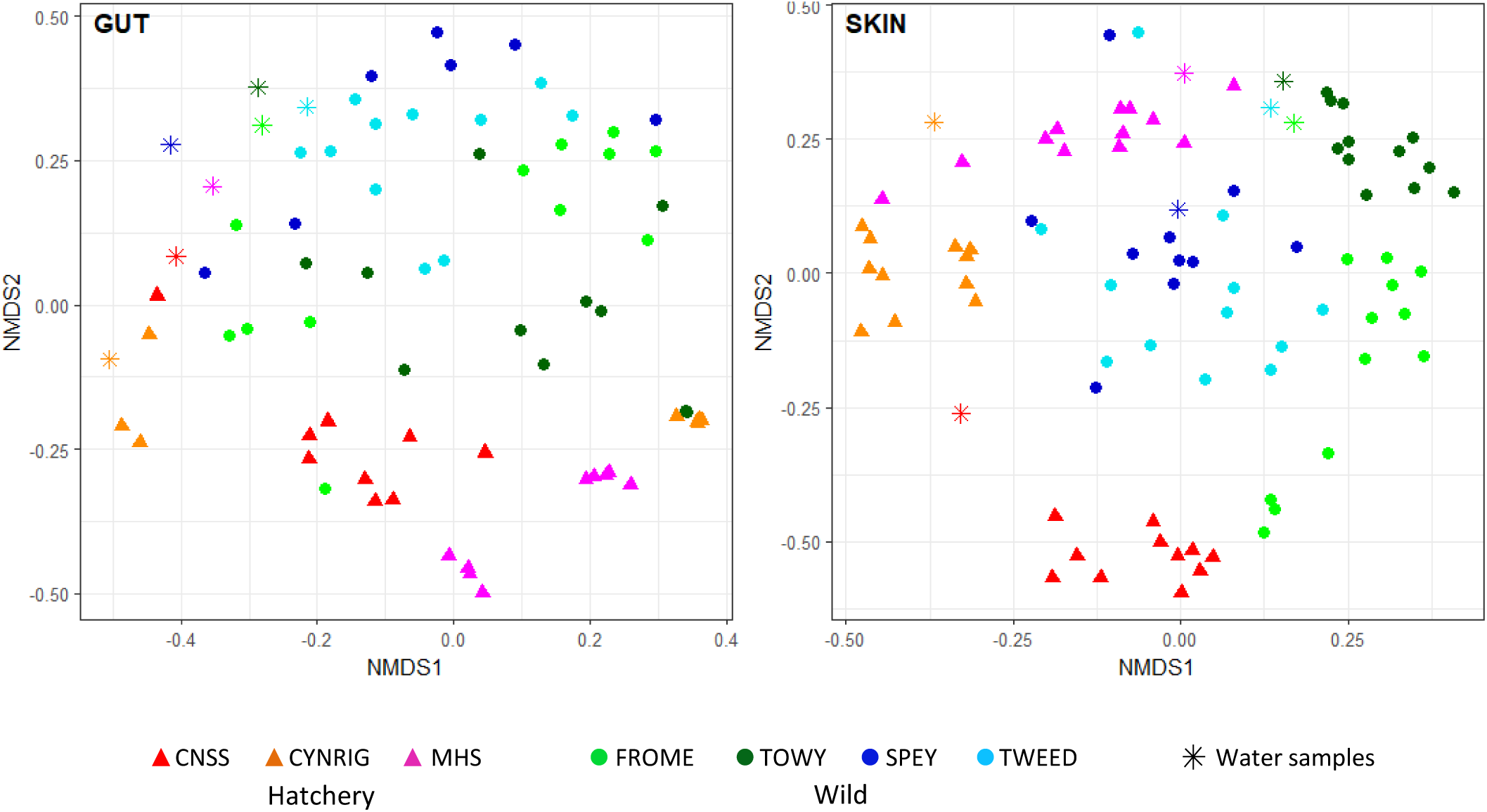
Non-metric multidimensional structure clustering of microbial community structure based on weighted Unifrac distances. Triangles represent hatchery individuals, circles represent wild fish, and stars represent water samples.

To investigate to what extent the water community structure influenced fish skin and gut microbial communities, average weighted Unifrac distances between each individual and the site water samples were calculated for each population. There was a significantly greater distance between the gut and water than the skin and the water (t_12_=3.69, *P=*0.003); average gut-water distance was 0.55 ±0.03 and the average skin-water distance was 0.43 ±0.02.

### Microbial diversity in skin, gut and water samples

Analysis of microbial diversity was performed at the OTU level, based on 97% sequence similarity. A total of 4,473 OTUs were identified in the salmon skin and 2,895 were identified in the intestine, 1,270 of which were shared between tissues. Approximately 27% of the OTUs found in the skin (1,209) and 30% of those found in the gut (877 OTUs) were also present in the water samples, and there were an additional 1,224 OTUs only found in water. Microbial diversity differed significantly between tissues and water (Chao1, *F*_2,161_= 80.39, *P*< 0.001; Shannon, *F*_2,161_= 16.55, *P*< 0.001), with highest microbial α-diversity found in water samples, followed by the fish skin, and the gut (Figure 3), and all pairwise comparisons were statistically significant (Tukey HSD; Chao1: gut-water *P*_adj_< 0.001, skin-water *P*_adj_< 0.001; skin-gut *P*_adj_= 0.029; Shannon: gut-water *P*_adj_< 0.001, skin-water *P*_adj_= 0.0489; skin-gut *P*_adj_< 0.001). Analysis of paired tissue samples for individual fish indicates that the skin microbiome was 31-36% more diverse than that of the gut (Chao1 *t*_71_ = 4.236, *P*< 0.001; Shannon *t*_71_ = 5.160, *P*< 0.001).

**Figure 3.**
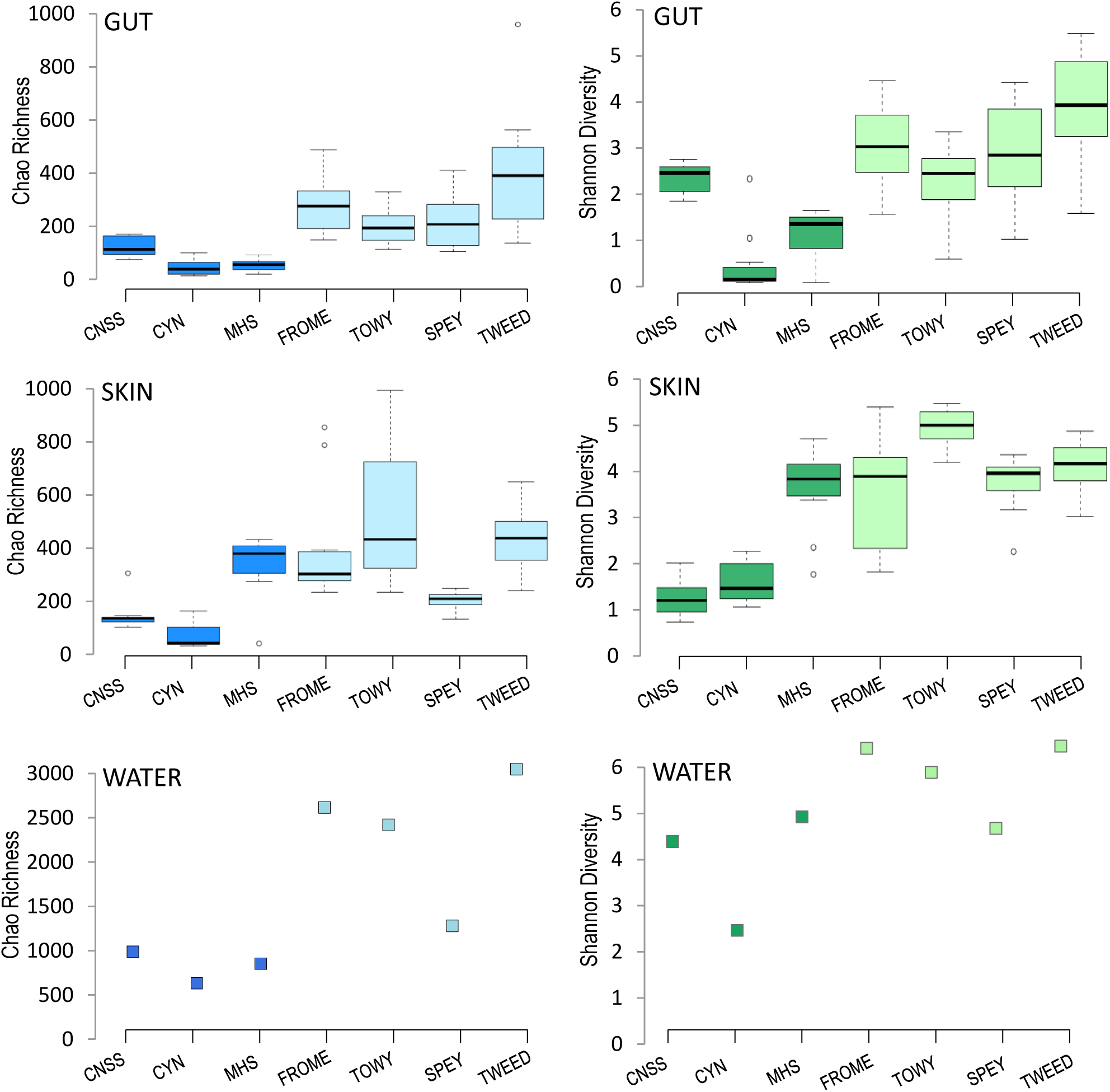
Measures of microbial α-diversity (Chao1 richness and Shannon diversity) in the gut and skin of juvenile Atlantic salmon (n = 10-12 fish/pop.), and in water samples (n=1 sample/pop.) at each site. Dark shaded bars represent hatchery populations and light shaded bars represent wild populations.

Water collected from rivers had c. 3 times higher microbial richness than water sampled in hatcheries (Chao1, *t*_3.44_= 3.876, *P*=0.023), although Shannon diversity indices were not statistically different (Shannon, *t*_3.22_= 2.258, *P*=0.103). Not surprisingly, water had a marked influence on the fish microbial diversity. At population level, a strong correlation in α-diversity was found between the water and the skin microbiome (Chao1 *r* = 0.818, *P*= 0.02; Shannon *r* = 0.720, *P*= 0.068), and also the gut (Chao1 *r* = 0.909, *P*= 0.004; Shannon *r* = 0.797, *P*= 0.032).

### Fish microbial diversity across populations

Overall, wild populations had considerably higher microbial diversity than hatchery populations, both in the skin (Chao1, *t*_78.43_= 5.29, *P*<0.001; Shannon, *t*_59.62_= 7.36, *P*< 0.001) and the gut (Chao1, *t*_52.39_= 8.24, *P*< 0.001; Shannon, *t*_71.20_= 7.31, *P*< 0.001; Figure 3). However, as this may have been due in part to the higher diversity of river water compared to hatchery water, we used water microbial diversity as an offset covariate. Microbial diversity continued to be significantly higher in wild salmon than in hatchery fish once the effect of water had been taken into account for both the gut (Chao1, *t*_4.998_= −3.214, *P*= 0.024; Shannon, *t*_4.86_= −2.289, *P*= 0.072) and the skin (Chao1, *t*_5.00_= −3.298, *P*= 0.021; Shannon, *t*_5.287_= 3.386, *P*= 0.018). Overall, variance component analysis indicated that 49-52% of variation in gut diversity was explained by whether the fish were of wild or hatchery origin, and 13-23% was explained by variation among populations. Results for skin diversity were similar: 28-52% of variation was explained by group origin and 29-31% was explained by population.

Other factors with a significant effect on microbial diversity in the model once the effects of group membership and water diversity were controlled for included body size, which was negatively associated with gut microbial diversity (Shannon, *t*_7.73_= −3.515, *P* = 0.008), and individual MHC heterozygosity, which had a marginal effect on skin diversity (Shannon, *t*_63.12_= −1.997, *P* = 0.050). No effect of sex, individual total heterozygosity, or condition factor was found on gut or skin microbial diversity (*P* > 0.5 in all tests).

Additionally, linear models were used to examine the effects of fish genetic diversity on microbial alpha diversity at the population level in addition to the individual-level analyses described above. No significant associations were found for heterozygosity, but populations with high allelic richness for neutral markers had higher microbial alpha diversity in the gut (Shannon; *R*^2^ = 0.68, *P* = 0.02; Chao1; *R*^2^ = 0.54, *P* = 0.06), but not in the skin (Shannon; *R*^2^ = 0.25, *P* = 0.14; Chao1; *R*^2^ = 0.07, *P* = 0.27). For the two MHC-linked markers combined, allelic richness had no effect on microbial diversity in the gut (Shannon; *R*^2^ = 0.52, *P* = 0.07; Chao1; *R*^2^ = 0.43, *P* = 0.11), but there was some evidence of a positive effect on the skin (Shannon; *R*^2^ = 0.67, *P* = 0.02; Chao1; *R*^2^ = 0.51, *P* = 0.07).

### Compositional analysis of microbial communities

In the gut, only one ‘core OTU’ (present in ≥ 80% of all individuals) was identified; *Pseudomonas* sp. However, when considering just wild fish, 12 additional core OTUs were identified, the majority of which were also Proteobacteria. In contrast, only one core gut OTU, *Mycoplasma* sp., was identified when considering all hatchery fish alone. For the skin, 13 OTUs were present in at least 80% of all individuals, all of which were also Proteobacteria.

Relative abundance of OTUs was notably distinct between populations, and between fish from hatchery or wild origin (Figure 4). A total of 130 OTUs showed significantly differential abundance (FDR <0.05) between hatchery and wild fish for the gut, the majority of which (100) were enriched in wild fish (Table S3). Of these, the largest proportion (44%) were Proteobacteria (predominantly Alphaproteobacteria), while over-represented gut OTUs in hatchery fish were dominated by Firmicutes (64%), of which the majority were Lactobacillales. For the skin, a total of 110 OTUs were differentially abundant between wild and hatchery fish, the majority of which (89) were also enriched in wild fish and were Proteobacteria (Table S4).

**Figure 4.**
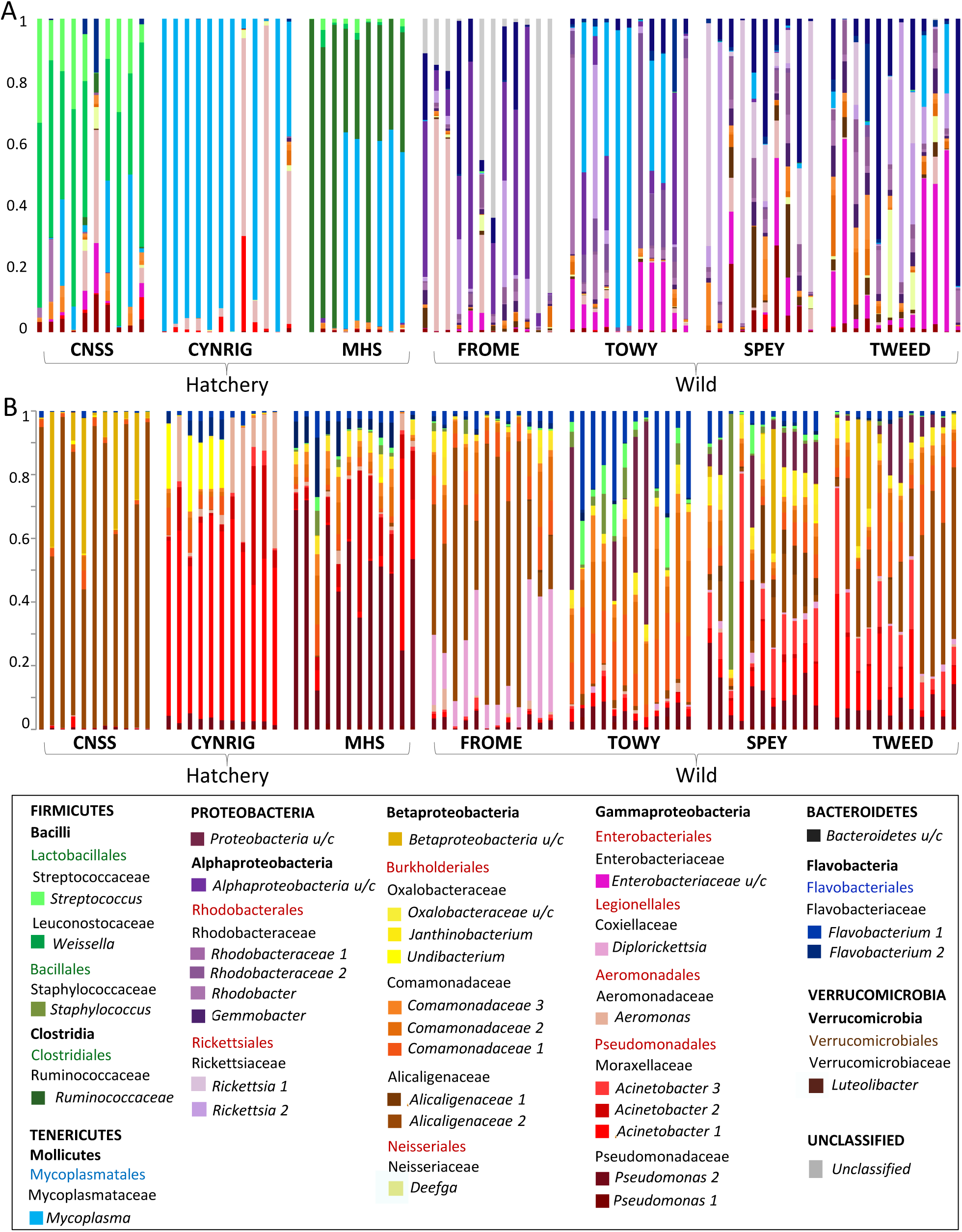
Relative abundance of the most abundant 35 OTUs across all samples, in (A) gut and (B) skin microbial communities. Each bar represents an individual fish.

Although distinct at the OTU level, skin and water samples had a similar phylum-level composition (Figure 5), and were strongly dominated by Proteobacteria, with lower abundance of Bacteroidetes, Firmicutes, Verrucimicrobia, Planctomycetes and Actinobacteria. The wild gut samples also showed a similar distribution of Phyla, although were dominated to a lesser extent by Proteobacteria and had a higher abundance of unclassified bacteria. In contrast, the hatchery gut samples comprised an entirely separate cluster, completely distinct from those found in the skin, water and wild gut samples, with elevated abundance of Firmicutes and/or Tenericutes which were less common – or even absent – among wild fish.

**Figure 5.**
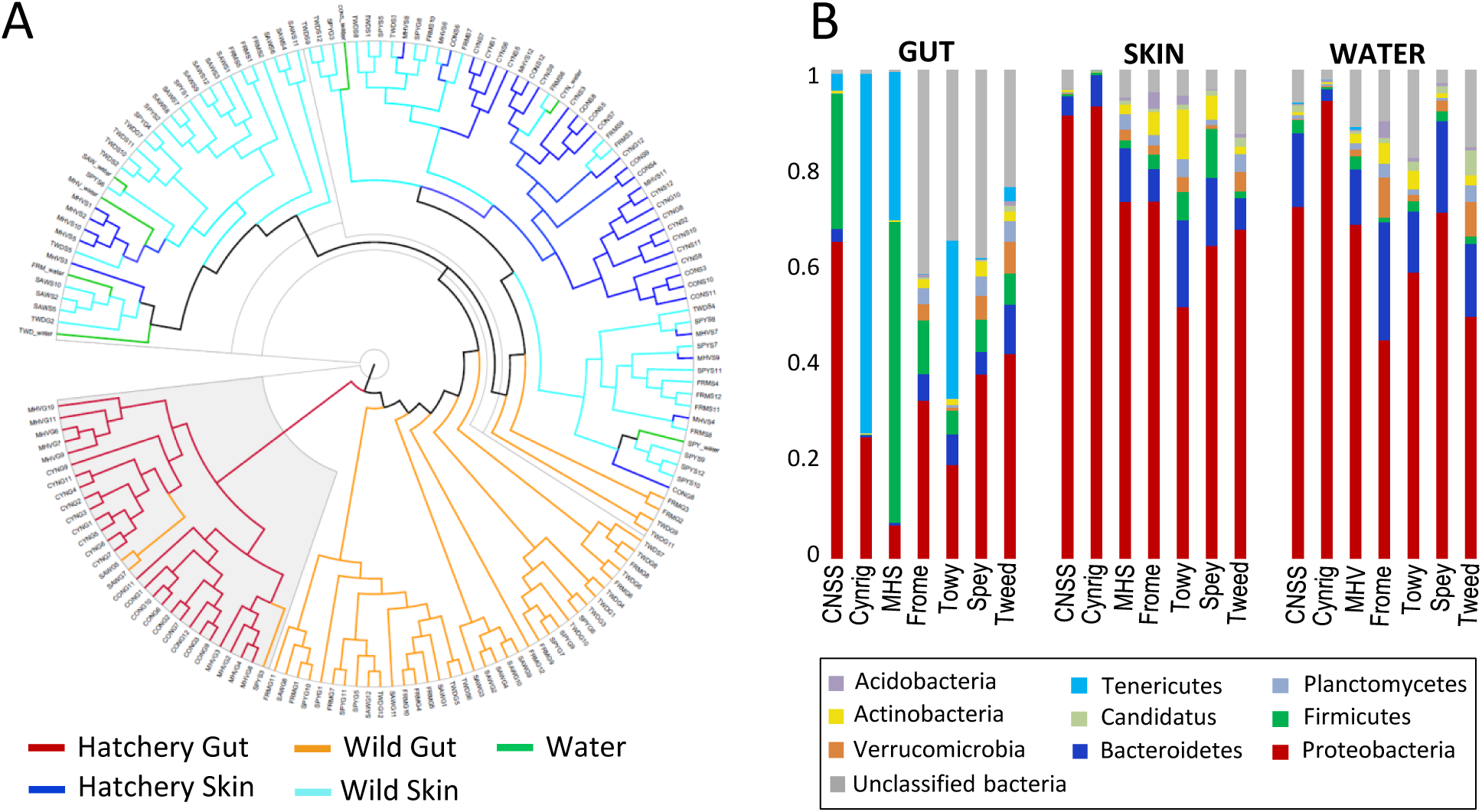
Phylum level analysis of microbial community structure; (A) cluster analysis based on Yue and Clayton measure of dissimilarity for all individual samples; (B) relative phyla abundance for each population.

### Functional analysis

Euclidean clustering of gut microbial communities based on metabolic function, broadly separated fish based on wild or hatchery origin, but there was no clear clustering of individual populations. Seven metabolic functions were significantly differentially represented between wild and hatchery-reared groups (FDR <0.05); ‘xylan degrader’, ‘sulphide oxidiser’, ‘sulphate reducer’, ‘iron reducer’ and ‘ammonia oxidiser’ were enriched in wild fish, while ‘nitrogen fixation’ and ‘denitrifying’ were enriched in hatchery fish (Figure 6a). In contrast, for the skin there was evidence of clear separation of metabolic function between individual populations, but there was no overall separation of wild and hatchery fish, although the functional terms ‘lignin degrader’ and ‘naphthalene degrading’ were significantly enriched in wild fish (Figure 6b).

**Figure 6.**
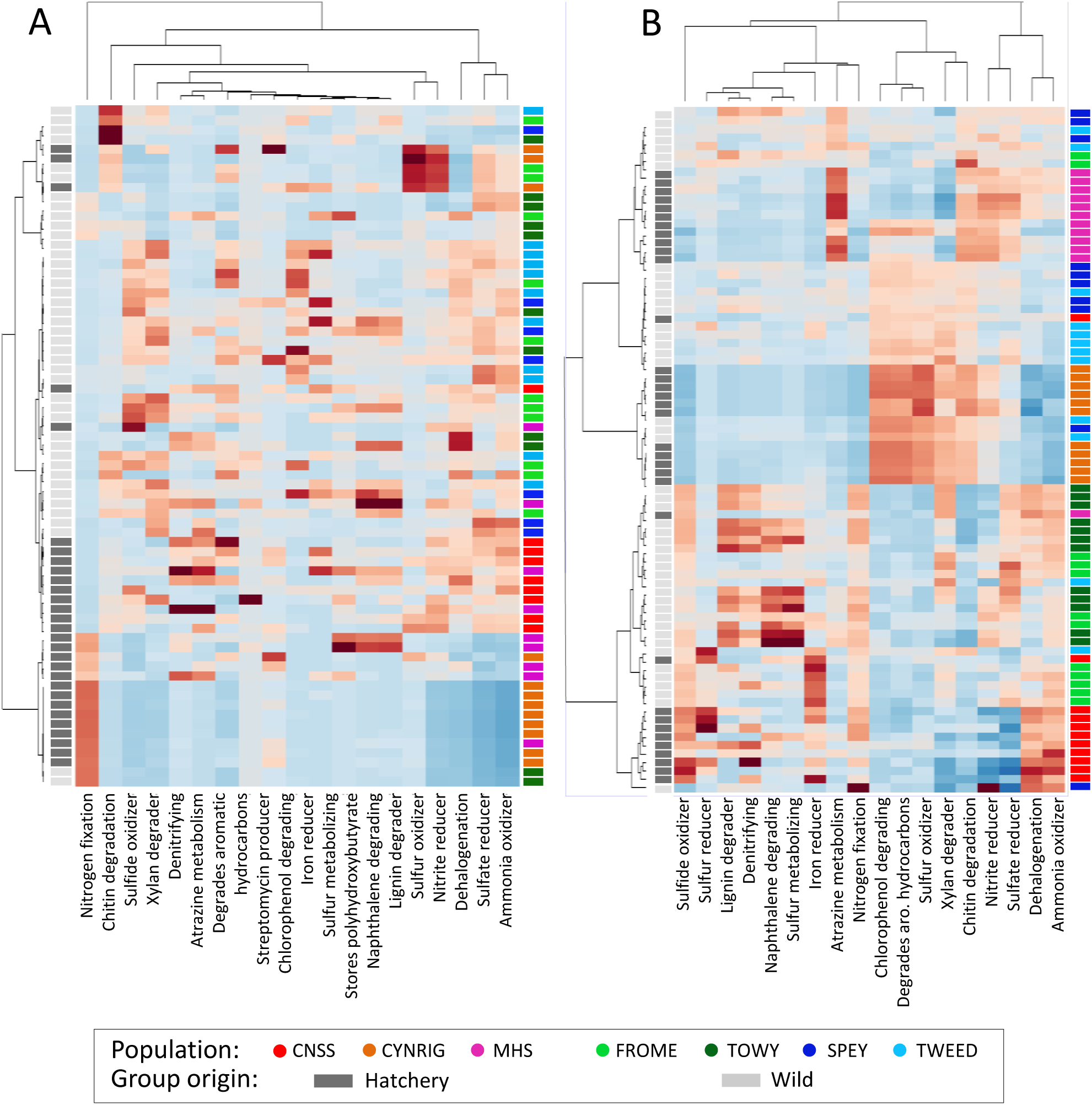
Functional analysis of microbial communities in (A) gut and (B) skin. OTU phenotype mapping was performed using METAGENassist, followed by functional analysis of community structure, based on ‘metabolism’, using Euclidean distance clustering.

## Discussion

Our study indicates that Atlantic salmon gut and skin microbial communities differ substantially, not only between fish living in the wild and in captivity, but also among populations. Surrounding water appeared to influence the diversity and structure of microbial communities of both the skin and the gut, but most of the differences identified among populations could not be explained by the effect of water alone. We identified a fundamental difference in the diversity, structure and function of intestinal microbial communities of wild and hatchery-reared fish, which likely reflect contrasting diets. We also found that host genetic variation was associated with diversity and structure of the gut and skin microbiome.

### A core microbiome?

It has been suggested that core microbiota have key functional importance in the symbiotic community assembly, and are present in the majority of individuals (Roeselers et al. 2011). A number of studies have reported a core gut microbiome for a range of fish species (Gajardo et al. 2016; Lowrey et al. 2015). However, the extent to which core microbiota are present across the range of a species is unclear. In our study we only detected one OTU (*Pseudomonas* sp.) in the gut of >80% individuals, present at low levels, suggesting a lack of species-wide core gut microbiota for Atlantic salmon fry across natural and captive populations. However, across wild populations there was evidence of a more extensive core gut microbiome (13 OTUs), composed predominantly of Proteobacteria OTUs, a number of which were amongst the most abundant in wild fish, suggesting a crucial function in natural salmon populations. Yet, these microbiota, which included several unclassified bacteria and OTUs from the Rhodobacteraceae, Rickettsiaceae, Enterobacteriaceae and Comamonadaceae families, were rare or entirely absent in captivity. These results argue against the existence of obligate core gut microbiota for this species when including the differential selective pressures of natural environments and captivity. Furthermore, the observed differences in associated metabolic processes between wild and hatchery-reared salmon suggest broader changes in community function.

As expected, the microbial communities present on salmon skin were very different from those found in the intestine, and were 31-36% more diverse, likely reflecting different functional roles (Lowrey et al. 2015). In contrast to the gut, we also found a more extensive core skin microbiome, with 11 OTUs (predominantly Proteobacteria) present in >80% of fish across all natural and captive populations, thus suggesting that deterministic factors shape the microbiome in a tissue-specific manner.

### Microbiome variation between populations

Despite this evidence for a core skin microbiome for Atlantic salmon, as well as core gut microbiota for wild fish, the relative abundance of shared OTUs was variable between populations. In fact, based on weighted Unifrac distances, all populations had distinct microbial communities in the gut and, especially, on the skin, likely reflecting differential environmental and host-specific filtering.

In particular, there were very marked differences in gut microbial diversity and structure between wild and hatchery-reared fish. Structural differences were apparent not just at the OTU level, but all the way up to Phylum level. Compared to the water, skin, and wild gut communities, hatchery gut communities were composed of a far more limited number of phyla, with an increased abundance of Firmicutes and Tenericutes. Together with the loss of core gut microbiota found in natural populations, and the observed distinction in metabolic processes, this suggests a fundamental change in the structure and function of the gut microbial communities of captive salmon. Hatchery fish also had lower microbial diversity, as found previously for mummichog (Givens et al. 2015) and for Atlantic salmon kept in a semi-natural environment (Dehler et al. 2017). These changes reflect the pronounced differences in rearing conditions; hatchery fish were fed an artificial diet, lived in waters with impoverished water microbial communities, and were also larger and had lower genetic diversity than wild fish.

Compared to the gut, there was less differentiation in skin microbiome structure and function between wild and hatchery fish, although there was a notable increase in the abundance of the order Pseudomonadales (primarily *Actinobacter* sp. and *Pseudomonas* sp.) in many of the hatchery fish, which have been associated with stressful conditions in salmonids (Boutin et al. 2013).

### Deterministic factors contributing to microbiome variation

In contrast to the extensive variation in microbiome structure observed among populations, we found a relatively high degree of convergence in microbial community structure among individuals within populations, both for the gut and the skin. Notably, captive populations had particularly low levels of inter-individual variation, likely reflecting more homogenous environmental conditions, a greater degree of inter-host microbial dispersal and lower genetic diversity. This suggests that population-specific deterministic factors shape fish microbial community structure.

The microbial communities present in the water are thought to determine the initial colonisation of the fish microbiota via direct seeding and by promoting the colonisation of other species (Giatsis et al. 2015; Ingerslev et al. 2014; Smith et al. 2015). We found that microbial diversity was highest in water samples (mean effective number of species, ENS = 189), followed by the fish skin (ENS = 49), and the gut (ENS = 16), with c. 30% of the OTUs identified in the fish gut and the skin were also present in the water. The lower microbial richness of hatchery water samples, reflecting the effects of water treatment in aquaculture, likely contributed to the reduced gut and skin microbial richness, and the lower degree of inter-individual variation in microbial structure, observed in captive fish. However, our results indicate that most of the differences in fish microbial diversity among populations remain, even when the effects of water are partialled out. Water microbial community structure was more similar to salmon skin than the gut, based on weighted Unifrac distances, number of shared OTUs and relative abundance of phyla. Nevertheless, both the skin and gut microbiomes were clearly distinct from the water samples. For example, the most abundant OTU in the skin, from the Alcaligenaceae family, was rare in the water, while the most abundant gut OTUs, including *Mycoplasma* sp., were absent in all water samples. This indicates that the fish gut and skin have specialised microbial communities that are influenced by, but remain distinct, from the free-living microbial communities present in the surrounding environment, as shown for other species (Larsen et al. 2013; Sullam et al. 2012).

Diet is likely to be the dominant factor shaping the microbial community of the fish gut. In sticklebacks, diet appears to have a greater influence on the gut microbiome than the surrounding water (Smith et al. 2015), and has a major influence on the gut of cultured salmonids (Llewellyn et al. 2014). In the wild, juvenile Atlantic salmon typically feed on a rich diet consisting of a large number of different aquatic and terrestrial macro-invertebrates (Orlov et al. 2006), which is more diverse and variable, both temporally and spatially, than artificial hatchery diets. The three hatchery populations were each fed a different commercial feed, consisting of different proportions of fish meal, fish oil and vegetable proteins. The increased abundance of Firmicutes observed in two of the hatcheries in this study is consistent with the use of plant-based fish feeds, as previously reported for other farmed salmonids (Ingerslev et al. 2014; Reveco et al. 2014). Differences in dietary composition therefore likely explain the differences found in the gut microbial communities of hatchery and wild salmon, including the absence of core OTUs across all salmon populations. In humans, culturally-driven dietary changes have similarly radically altered gut microbial community structure, the symbiotic balance between host immune system and the microbiota, and the evolution of this association (Walter & Ley 2011).

Mammalian studies have highlighted how host genetic background can influence gut microbial communities via the immune system and complex metabolic pathways (Blekhman et al. 2015), but less is known for other taxa. For example, in order to prevent inappropriate immune responses, mammalian hosts appear to distinguish between beneficial mutualists and harmful pathogens, and individuals displaying high immunocompetence tend to display high gut microbial diversity (Kamada et al. 2013). In stickleback, high MHC variation has been associated with a diverse gut microbiota (Bolnick et al. 2014), and our study also found some evidence for a role of host genetic diversity on the salmon microbiome. Population-level allelic richness was positively associated with high gut microbial diversity, while variation in the skin microbiome was linked with variation for two MHC-linked markers. There was also a positive relationship between genetic distance and community Unifrac distance for all individuals for both the gut, and to a lesser extent, the skin. We also observed that the two natural populations (Spey and Tweed) that were most genetically similar had the most similar gut microbial communities, as previously seen in sticklebacks (Smith et al. 2015) and the Trinidadian guppy (Sullam et al. 2015).

### Conclusions and perspective

As we predicted, we found evidence of considerable variation in the gut and skin microbiome between populations of juvenile Atlantic salmon and relatively high convergence within populations, especially for captive fish. These differences seemingly reflect local variation in water, in diet, and in genetic diversity. As expected, these deterministic factors influencing variation in the skin and gut microbiome appeared to act in a tissue-specific manner. Between natural and hatchery-reared fish we found evidence of a fundamental difference in the diversity, structure and function of microbial communities in the intestine, but not on the skin, likely reflecting contrasting selective pressures in captivity and in the wild. Reduced microbial diversity has generally been associated with stress and ill health (Hooper et al. 2012), and Rosshart (2017) recently reported that evolution of the laboratory mouse microbiome in restricted conditions has induced significant fitness costs related to immunity. On the other hand, it is possible that microbial community specialisation may contribute to the process of fish domestication, and that microbiome plasticity may even be harnessed to enhance certain phenotypic traits and improve the health and fitness of fish in aquaculture. Microbial communities can have a vital influence on host phenotype, and may contribute to host plasticity, acclimation to environmental change or local adaptation of natural populations. However, variation in microbiome communities has rarely been linked to differences in host fitness for any species, and this is a priority for future research.

## Acknowledgements

We are grateful to Deiene Rodriquez-Barreto and Chloe Robinson for assistance with sampling, Rasmus Lauridsen, Ronald Campbell, Mark Coulson, Brian Shaw, and Stuart Rees for sampling of wild salmon, and Patrick Martin (CNSS), Dave Cockerill (Marine Harvest Scotland) and John Taylor (NRW Cynrig Hatchery) for providing us with the hatchery samples. This work was funded by a BBSRC-NERC Aquaculture grant (BB/M026469/1) to GGL and the Welsh Government and Higher Education Funding Council for Wales (HEFCW) through the Sêr Cymru National Research Network for Low Carbon Energy and Environment (NRN-LCEE) to SC.

## Data Accessibility

All raw sequence reads have been deposited in the European Nucleotide Archive under study accession number PRJEB22688.

## Author contributions

TUW, SC and CGL designed research, TUW and MH performed research, TUW analysed data and drafted the manuscript. All authors contributed to and approved the final version of the manuscript.

