## Supplementary Materials for "Inter-population variation in the Atlantic salmon microbiome reflects environmental and genetic diversity"

**This supporting information contains:**

**Page 2-4:** Supplementary methods; population genetics analysis, 16S rRNA sequencing and bioinformatics analysis

**Page 5:** Figure S1. Principal coordinate analysis based on Nei's pairwise genetic distance between individual fish. Triangles represent hatchery fish while circles represent wild fish.

**Page 6:** Table S1. Measures of genetic diversity in the study salmon populations at 12 neutral and two MHC-linked microsatellite DNA loci.

**Page 7:** Table S2. Pairwise genetic distances (above diagonal) and $F_{ST}$ (below diagonal) between study populations screened at 14 microsatellite DNA loci (*$P$ < 0.01).

**Page 8-12:** Table S3. Differentially abundant OTUs identified between wild and hatchery fish in the gut.

**Page 13-16:** Table S4. Differentially abundant OTUs identified between wild and hatchery fish in the skin.

**Page 17:** References

**Supplementary methods**

*Population genetics analysis*

DNA loci were amplified using the Qiagen Multiplex PCR mastermix in two multiplex reactions with final primer concentrations as follows; a) 0.05 µM *SSsp2210*, 0.2 µM *SSspG7*, 0.125 µM *SSaD144*, 0.1 µM *SSa202*, 0.15 µM *SSsp2201*, 0.15 µM *SSsp1605*, 0.05 µM *Sasa-UBA* and b) 0.25 µM *SSa197*, 0.15 µM *Ssosl85*, 0.18 µM *SSsp3016*, 0.1 µM *SSa171*, 0.23 µM *SSsp2216*, 1.12 µM *SSa289*, 0.61 µM *Sasa-DAA.* Each reaction was performed in a total volume of 8 µl, using 2 µl of genomic DNA extracted from the gut. PCR conditions consisted of 95 °C for 15 min; followed by 8 cycles of 94 °C for 30s, 64>56 °C touchdown for 90s, 72 °C for 90s; then 24 cycles of 94 °C for 30s, 56 °C 90s, 72 °C for 90s; then a final extension 72 °C for 10 mins.

*DNA extraction and 16S rRNA sequencing*

DNA extraction from all gut and skin swab samples was performed using MoBio PowerSoil® DNA Isolation Kit (Cambio, Cambridge, UK) according to the manufacturer's instructions, with an additional incubation step of 10 minutes at 65 °C prior to bead beating. Water samples were centrifuged at 5000xg for 1 hour at 4°C before DNA was extracted from the pellet. DNA concentration and purity was assessed using a NanoDrop ND-1000 Spectrophotometer (NanoDrop Technologies, Wilmington, USA). 16S library preparation using Nextera XT Index kit was performed according the Illumina Metagenomic Sequencing Library Preparation guide using 12.5 ng total genomic DNA. The V4 hypervariable region of the bacterial 16S gene was amplified using the primers selected as the best candidates for bacterial and archaeal representation (Klindworth et al. 2013); 519F (5'-CAGCMGCCGCGGTAA) and 785R (5'-TACNVGGGTATCTAATCC), each with 5' tags for barcode attachment. Reaction conditions for the first PCR amplification consisted of an initial denaturation at 95°C for 3 min, followed by 25 cycles of 95°C for 30s, 55°C for 30s and 72°C for 30s, then a final elongation at 72°C for 5 min, using 12.5 ng genomic DNA, 0.2 µM of primers and KAPA HiFi HotStart ReadyMix (Kapa Biosystems, London, UK) in a total volume of 25 µl. All products were purified with Agencourt Ampure XP beads (Beckman Coulter then used as template for the second PCR reaction to add indexed sequencing adaptors to each library (Nextera XT Indices, Illumina). The reaction conditions used were the same as before, but using eight cycles, and a total reaction volume of 50 µl. The final product (420 bp) was verified and size selected from a 2 % agarose gel, and purified with

Ampure XP beads. All samples were quantified using a Qubit 3.0 Fluorometer (Thermo Fisher Scientific, UK) and pooled in equal concentrations, before sequencing using an Illumina MiSeq (300 bp PE reads).

*16S rRNA bioinformatics analysis*

Adaptor contamination and poor quality bases from the 3' end were removed from the raw sequence reads using a sliding window of 4 bp and a minimum quality score of Q=20 in Trimmomatic (Bolger et al. 2014). Microbial community analysis was then conducted using mother v1.37 (Kozich et al. 2013), Qiiime v1.9 (Caporaso et al. 2010) and R v3.3.2 (R_Core_Team 2014). Forward and reverse reads were merged and then filtered to retain amplicons of the target size range (260-300 bp) and remove those containing ambiguous bases. Contigs were aligned to the Silva seed reference database (version 123) (Quast et al. 2013) which was first trimmed to include only the target V4 region. For contigs with high quality alignments, further noise reduction was performed using mothur's pre-clustering algorithm. Potential chimeras were then removed through implementation of UCHIME (Edgar et al. 2011) in mothur before taxonomic classification using the Silva reference taxonomy and removal of mitochondrial, eukaryote and chloroplast sequences.

In order to examine microbial community structure, phylogenetic trees were constructed using all retained sequences, using the relaxed neighbour joining method implemented in Clearcut (Evans et al. 2006), within mothur. Weighted Unifrac (Lozupone et al. 2011) distances between samples, incorporating sequence phylogeny and relative abundance, were then calculated and used for analysis of microbial community structure. Non-metric multidimensional scaling analysis was performed separately for gut and skin samples in mothur. After checking that data met assumptions of homogeneity of variance using Betadisper, statistical analysis of Unifrac distances was performed using Adonis (both within the Vegan package for R; (Oksanen et al. 2017)). We assessed effects of origin (wild/hatchery), population, fork length, condition factor, sex, individual heterozygosity and individual MHC heterozygosity on gut and skin structural community variation, and used the strata function to specify a nested model of population within in group origin. Additionally, HOMOVA (Stewart & Excoffier 1996), within mothur, was used to specifically quantify the degree of intra-population variation in community structure, and test for statistical differences in this degree of variance between populations. To investigate a hypothesised influence of fish genetic background on microbial community structure a Mantel test, using the Pearson method, was employed in mothur to test for correlation between individual level genetic distances and weighted Unifrac distances for both the gut and the skin.

Analysis of microbial community alpha diversity and composition were then performed at the operational taxonomic unit (OTU) level, following clustering of sequences in mothur based on 97% sequence similarity. Several samples had low numbers of good quality microbial sequence reads. In order to maximise sample inclusion whilst ensuring high Good's coverage (≥ 94%) for all included samples, reads were subsampled to a depth of 4012/sample. Seventy six gut and 81 skin samples (min of 10 samples per population), together with a water sample from each site, were retained and used for analysis of alpha diversity. We calculated two measures of alpha diversity (Chao1 richness and Shannon diversity) to examine variation in microbial diversity using mothur. For illustrative purposes, we also converted the Shannon index to an Effective Number of Species (ENS, i.e. the number of equally abundant species in an ideal even community) as per Jost (Jost 2006).

We analysed variation in alpha diversity by linear mixed modelling with the *lme4* package in R using group (hatchery vs. wild), fork length, condition factor, sex, individual heterozygosity and individual MHC heterozygosity as fixed factors, and population as a random factor to account for spatial autocorrelation. Given the influential role of water on the fish microbiome, we included the microbiome diversity of the water samples at each site as an *offset* to statistically control for the effects of the surrounding water on fish microbial diversity. Model simplification was achieved by single deletion tests using the *drop1* command via maximum likelihood; the minimal adequate model was then refitted by restricted maximum likelihood(Zuur et al. 2009), and the Satterthwaite approximation was used to obtain approximate significance levels using the *lmerTest* package. Variance components were calculated using the *VarCorr* function.

OTUs that were present in at least 80% of all individuals, as well as separately in 80% of wild and in 80% of hatchery fish, were identified for the gut and for the skin samples using the compute_core_microbiome function in Qiime. Following filtering of singleton OTUs from the dataset, a Kruskal-Wallace test, incorporating FDR calculation for multiple testing, was implemented using the group_significance function in Qiime to identify skin and gut OTUs with significantly differential abundance between fish from a wild and hatchery origin. In addition to these OTU-level analyses further community composition analysis was performed at the Phylum level using the summarize_taxa function in Qiime, followed by cluster analysis of all gut, skin and water samples using the Bray-Curtis similarity index and visualisation using FigTree v1.4.3 (Rambaut 2007).

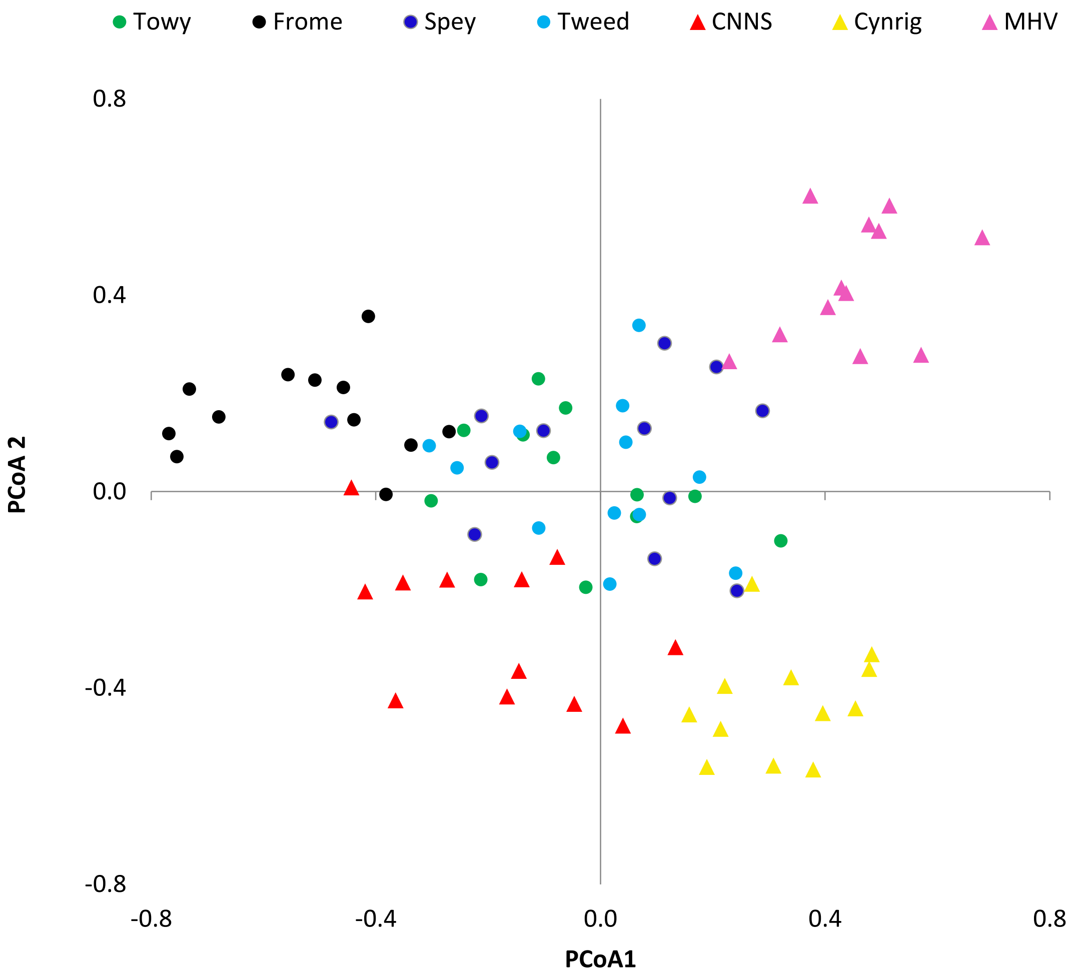


**Figure S1.** Principal coordinate analysis based on Nei's pairwise genetic distance between all individual fish. Triangles represent hatchery populations while circles represent wild fish.

**Table S1.** Measures of genetic diversity in the study salmon populations at 12 neutral and two MHC-linked microsatellite DNA loci.

| **Type/Population** | *N* | **Observed Heterozygosity** | | **Allelic richness** | |
|---|---|---|---|---|---|
| | | neutral | MHC | neutral | MHC |
| **Hatchery** | | | | | |
| CNSS | 12 | 0.73 | 0.57 | 5.90 | 4.49 |
| CYN | 12 | 0.86 | 0.68 | 3.48 | 3.00 |
| MHS | 12 | 0.79 | 0.46 | 5.82 | 5.11 |
| **Wild** | | | | | |
| R. Towy | 12 | 0.90 | 0.71 | 6.45 | 6.05 |
| R. Frome | 12 | 0.73 | 0.75 | 5.86 | 4.82 |
| R. Spey | 12 | 0.87 | 0.83 | 8.43 | 6.09 |
| R. Tweed | 12 | 0.81 | 0.83 | 8.29 | 6.14 |

**Table S2.** Pairwise genetic distances (above diagonal) and $F_{ST}$ (below diagonal) between study populations screened at 14 microsatellite DNA loci (*$P$ < 0.01).

| | **Hatchery** | | | **Wild** | | | |
|---|---|---|---|---|---|---|---|
| | CNSS | CYN | MHS | Frome | Towy | Spey | Tweed |
| CNSS | | 28.99 | 31.63 | 29.84 | 28.72 | 28.73 | 29.67 |
| CYN | 0.14* | | 27.68 | 29.15 | 27.00 | 26.29 | 26.69 |
| MHS | 0.16* | 0.15* | | 30.83 | 28.22 | 26.86 | 28.17 |
| Frome | 0.13* | 0.20* | 0.21* | | 27.01 | 26.39 | 27.68 |
| Towy | 0.09* | 0.13* | 0.12* | 0.12* | | 25.06 | 27.01 |
| Spey | 0.08* | 0.10* | 0.09* | 0.10* | 0.04* | | 25.27 |
| Tweed | 0.07* | 0.08* | 0.10* | 0.10* | 0.05* | 0.01 | |

**Table S3.** Differentially abundant OTUs identified between wild and hatchery fish in the gut.

| OTU | Test-Statistic | P | FDR_P | Wild mean abundance | Hatchery mean abundance | Direction | Phylum | Class | Order | Family | Genus |
|---|---|---|---|---|---|---|---|---|---|---|---|
| Otu000026 | 45.3 | 1.71E-11 | 1.06E-07 | 24.9 | 1.4 | W > H | Proteobacteria | Alphaproteobacteria | Rhodobacterales | Rhodobacteraceae | Rhodobacter |
| Otu000041 | 42.3 | 7.86E-11 | 1.63E-07 | 14.8 | 0.2 | W > H | Verrucomicrobia | Verrucomicrobiae | Verrucomicrobiales | Verrucomicrobiaceae | Luteolibacter |
| Otu000034 | 42.9 | 5.90E-11 | 1.63E-07 | 20.5 | 1.1 | W > H | Proteobacteria | Alphaproteobacteria | Rhodobacterales | Rhodobacteraceae | Rhodobacter |
| Otu000021 | 39.6 | 3.11E-10 | 3.22E-07 | 142.3 | 0.2 | W > H | Proteobacteria | Alphaproteobacteria | Rickettsiales | Rickettsiaceae | Rickettsia |
| Otu000093 | 39.7 | 2.89E-10 | 3.22E-07 | 17.9 | 0.0 | W > H | unclassified | Unclassified | unclassified | unclassified | unclassified |
| Otu000087 | 39.9 | 2.69E-10 | 3.22E-07 | 7.3 | 0.1 | W > H | Proteobacteria | Alphaproteobacteria | Rhizobiales | unclassified | unclassified |
| Otu000043 | 35.7 | 2.30E-09 | 2.05E-06 | 9.2 | 0.1 | W > H | Proteobacteria | Gammaproteobacteria | Alteromonadales | OM60 | unclassified |
| Otu000065 | 34.9 | 3.50E-09 | 2.72E-06 | 7.5 | 0.1 | W > H | Proteobacteria | Alphaproteobacteria | Rhodobacterales | Rhodobacteraceae | Rhodobacter |
| Otu000038 | 34.2 | 4.92E-09 | 3.40E-06 | 7.1 | 0.1 | W > H | Proteobacteria | Betaproteobacteria | Burkholderiales | Comamonadaceae | unclassified |
| Otu000205 | 34.0 | 5.46E-09 | 3.40E-06 | 5.0 | 0.0 | W > H | Firmicutes | Clostridia | Clostridiales | Clostridiaceae | SMB53 |
| Otu000136 | 32.4 | 1.28E-08 | 6.70E-06 | 5.5 | 0.0 | W > H | Verrucomicrobia | Verrucomicrobiae | Verrucomicrobiales | Verrucomicrobiaceae | unclassified |
| Otu000031 | 32.3 | 1.29E-08 | 6.70E-06 | 10.8 | 0.0 | W > H | Proteobacteria | Gammaproteobacteria | Xanthomonadales | Xanthomonadaceae | unclassified |
| Otu000068 | 30.4 | 3.60E-08 | 1.72E-05 | 17.7 | 0.0 | W > H | Cyanobacteria | Oscillatoriophycideae | Chroococcales | Xenococcaceae | Chroococcidiopsis |
| Otu000134 | 29.2 | 6.54E-08 | 2.88E-05 | 4.7 | 0.0 | W > H | Verrucomicrobia | Spartobacteria | Chthoniobacterales | Chthoniobacteraceae | Chthoniobacter |
| Otu000020 | 29.0 | 7.22E-08 | 2.88E-05 | 105.7 | 11.8 | W > H | Proteobacteria | Gammaproteobacteria | Enterobacteriales | Enterobacteriaceae | unclassified |
| Otu000010 | 29.0 | 7.41E-08 | 2.88E-05 | 18.4 | 4.2 | W > H | Proteobacteria | Betaproteobacteria | Burkholderiales | Comamonadaceae | Rhodoferax |
| Otu000128 | 27.7 | 1.44E-07 | 5.26E-05 | 19.7 | 0.0 | W > H | Proteobacteria | unclassified | unclassified | unclassified | unclassified |
| Otu000079 | 26.2 | 3.00E-07 | 9.57E-05 | 3.6 | 0.0 | W > H | Proteobacteria | Alphaproteobacteria | Sphingomonadales | Sphingomonadaceae | Novosphingobium |
| Otu000148 | 26.2 | 3.06E-07 | 9.57E-05 | 4.4 | 0.0 | W > H | Verrucomicrobia | Verrucomicrobiae | Verrucomicrobiales | Verrucomicrobiaceae | Luteolibacter |
| Otu000131 | 26.2 | 3.08E-07 | 9.57E-05 | 2.9 | 0.0 | W > H | Proteobacteria | Betaproteobacteria | unclassified | unclassified | unclassified |
| Otu000311 | 24.8 | 6.40E-07 | 1.89E-04 | 2.5 | 0.0 | W > H | Proteobacteria | Alphaproteobacteria | Rhodospirillales | Acetobacteraceae | Roseomonas |
| Otu000052 | 24.4 | 7.93E-07 | 2.24E-04 | 3.5 | 0.0 | W > H | Proteobacteria | Betaproteobacteria | Burkholderiales | Comamonadaceae | unclassified |
| Otu000297 | 23.5 | 1.27E-06 | 3.37E-04 | 2.4 | 0.0 | W > H | Actinobacteria | Actinobacteria | Actinomycetales | unclassified | unclassified |
| Otu000108 | 23.4 | 1.34E-06 | 3.37E-04 | 2.9 | 0.0 | W > H | Planctomycetes | Planctomycetia | Planctomycetales | Planctomycetaceae | Planctomyces |
| Otu000107 | 23.0 | 1.62E-06 | 3.47E-04 | 4.8 | 0.0 | W > H | Verrucomicrobia | Verrucomicrobiae | Verrucomicrobiales | Verrucomicrobiaceae | unclassified |

| | | | | | | | | | | | |
|---|---|---|---|---|---|---|---|---|---|---|---|
| Otu000191 | 22.2 | 2.42E-06 | 4.85E-04 | 4.3 | 0.4 | W > H | Firmicutes | Bacilli | Bacillales | Bacillaceae | Bacillus |
| Otu000054 | 22.0 | 2.74E-06 | 5.17E-04 | 5.2 | 0.0 | W > H | Proteobacteria | Betaproteobacteria | Methylophilales | Methylophilaceae | Methylotenera |
| Otu000157 | 22.0 | 2.74E-06 | 5.17E-04 | 3.3 | 0.0 | W > H | TM7 | TM7-1 | unclassified | unclassified | unclassified |
| Otu000540 | 20.8 | 5.06E-06 | 8.99E-04 | 1.3 | 0.0 | W > H | Firmicutes | Clostridia | Clostridiales | Clostridiaceae | Clostridium |
| Otu000331 | 20.7 | 5.29E-06 | 9.15E-04 | 3.0 | 0.0 | W > H | Planctomycetes | Planctomycetia | Gemmatales | Gemmataceae | unclassified |
| Otu000163 | 20.7 | 5.46E-06 | 9.17E-04 | 3.2 | 0.0 | W > H | Proteobacteria | Alphaproteobacteria | Rhizobiales | unclassified | unclassified |
| Otu000029 | 19.9 | 8.16E-06 | 1.34E-03 | 65.0 | 0.0 | W > H | unclassified | unclassified | unclassified | unclassified | unclassified |
| Otu000109 | 19.8 | 8.39E-06 | 1.34E-03 | 6.3 | 0.0 | W > H | Verrucomicrobia | Verrucomicrobiae | Verrucomicrobiales | Verrucomicrobiaceae | Luteolibacter |
| Otu000112 | 19.4 | 1.06E-05 | 1.48E-03 | 4.0 | 0.0 | W > H | Bacteroidetes | Cytophagia | Cytophagales | Cytophagaceae | Leadbetterella |
| Otu000113 | 19.4 | 1.07E-05 | 1.48E-03 | 2.6 | 0.0 | W > H | Proteobacteria | Gammaproteobacteria | Xanthomonadales | Sinobacteraceae | unclassified |
| Otu000258 | 19.4 | 1.07E-05 | 1.48E-03 | 3.7 | 0.0 | W > H | Proteobacteria | Betaproteobacteria | unclassified | unclassified | unclassified |
| Otu000178 | 19.4 | 1.07E-05 | 1.48E-03 | 3.2 | 0.0 | W > H | Planctomycetes | Planctomycetia | Planctomycetales | Planctomycetaceae | Planctomyces |
| Otu000147 | 19.4 | 1.07E-05 | 1.48E-03 | 3.3 | 0.0 | W > H | Verrucomicrobia | Verrucomicrobiae | Verrucomicrobiales | Verrucomicrobiaceae | unclassified |
| Otu000358 | 19.4 | 1.07E-05 | 1.48E-03 | 3.2 | 0.0 | W > H | Proteobacteria | Alphaproteobacteria | Sphingomonadales | Sphingomonadaceae | Zymomonas |
| Otu000472 | 18.2 | 2.02E-05 | 2.56E-03 | 1.3 | 0.0 | W > H | Actinobacteria | Actinobacteria | Actinomycetales | Nocardioidaceae | unclassified |
| Otu000150 | 18.2 | 2.03E-05 | 2.56E-03 | 2.2 | 0.0 | W > H | Bacteroidetes | Flavobacteriia | Flavobacteriales | Flavobacteriaceae | Flavobacterium |
| Otu000515 | 18.1 | 2.05E-05 | 2.56E-03 | 2.3 | 0.0 | W > H | Firmicutes | Clostridia | Clostridiales | Peptostreptococcaceae | unclassified |
| Otu000117 | 18.1 | 2.05E-05 | 2.56E-03 | 2.2 | 0.0 | W > H | Proteobacteria | Betaproteobacteria | Rhodocyclales | Rhodocyclaceae | Dechloromonas |
| Otu000143 | 18.1 | 2.06E-05 | 2.56E-03 | 2.1 | 0.0 | W > H | Proteobacteria | Gammaproteobacteria | Xanthomonadales | Sinobacteraceae | unclassified |
| Otu000144 | 17.3 | 3.24E-05 | 3.95E-03 | 6.0 | 0.1 | W > H | Bacteroidetes | Cytophagia | Cytophagales | Cytophagaceae | Emticicia |
| Otu000138 | 16.9 | 3.85E-05 | 4.13E-03 | 1.9 | 0.0 | W > H | Planctomycetes | Planctomycetia | Pirellulales | Pirellulaceae | unclassified |
| Otu000222 | 16.9 | 3.87E-05 | 4.13E-03 | 1.9 | 0.0 | W > H | Proteobacteria | Alphaproteobacteria | Rhizobiales | Phyllobacteriaceae | unclassified |
| Otu000198 | 16.9 | 3.87E-05 | 4.13E-03 | 1.7 | 0.0 | W > H | Planctomycetes | Planctomycetia | Gemmatales | Gemmataceae | unclassified |
| Otu000246 | 16.9 | 3.89E-05 | 4.13E-03 | 2.3 | 0.0 | W > H | Proteobacteria | Alphaproteobacteria | unclassified | unclassified | unclassified |
| Otu000186 | 16.9 | 3.89E-05 | 4.13E-03 | 2.6 | 0.0 | W > H | Planctomycetes | Planctomycetia | Pirellulales | Pirellulaceae | unclassified |
| Otu000248 | 16.9 | 3.90E-05 | 4.13E-03 | 3.8 | 0.0 | W > H | Proteobacteria | Betaproteobacteria | unclassified | unclassified | unclassified |
| Otu000221 | 16.9 | 3.90E-05 | 4.13E-03 | 2.6 | 0.0 | W > H | Verrucomicrobia | [Spartobacteria] | [Chthoniobacterales] | [Chthoniobacteraceae] | unclassified |
| Otu000069 | 16.9 | 3.91E-05 | 4.13E-03 | 16.0 | 0.0 | W > H | Bacteroidetes | Flavobacteriia | Flavobacteriales | [Weeksellaceae] | unclassified |
| Otu000249 | 16.6 | 4.53E-05 | 4.70E-03 | 4.7 | 0.1 | W > H | unclassified | unclassified | unclassified | unclassified | unclassified |
| Otu000045 | 16.3 | 5.27E-05 | 5.37E-03 | 5.1 | 0.9 | W > H | Bacteroidetes | Cytophagia | Cytophagales | Cytophagaceae | Flectobacillus |

| | | | | | | | | | | | |
|---|---|---|---|---|---|---|---|---|---|---|---|
| Otu000141 | 16.0 | 6.23E-05 | 6.15E-03 | 3.9 | 0.4 | W > H | Verrucomicrobia | Verrucomicrobiae | Verrucomicrobiales | Verrucomicrobiaceae | unclassified |
| Otu000598 | 15.8 | 7.00E-05 | 6.48E-03 | 1.4 | 0.0 | W > H | Proteobacteria | Alphaproteobacteria | Rhizobiales | Hyphomicrobiaceae | Hyphomicrobium |
| Otu000046 | 15.8 | 7.13E-05 | 6.48E-03 | 1.2 | 0.0 | W > H | Proteobacteria | Betaproteobacteria | Burkholderiales | Comamonadaceae | Hydrogenophaga |
| Otu000270 | 15.8 | 7.14E-05 | 6.48E-03 | 1.4 | 0.0 | W > H | Actinobacteria | Acidimicrobiia | Acidimicrobiales | C111 | unclassified |
| Otu000405 | 15.8 | 7.22E-05 | 6.48E-03 | 1.3 | 0.0 | W > H | Proteobacteria | Alphaproteobacteria | Rhizobiales | unclassified | unclassified |
| Otu000250 | 15.7 | 7.28E-05 | 6.48E-03 | 2.2 | 0.0 | W > H | unclassified | unclassified | unclassified | unclassified | unclassified |
| Otu000167 | 15.7 | 7.28E-05 | 6.48E-03 | 3.2 | 0.0 | W > H | Planctomycetes | Planctomycetia | Pirellulales | Pirellulaceae | unclassified |
| Otu000135 | 15.7 | 7.29E-05 | 6.48E-03 | 5.2 | 0.0 | W > H | Proteobacteria | Alphaproteobacteria | Rhizobiales | unclassified | unclassified |
| Otu000008 | 15.2 | 9.47E-05 | 7.96E-03 | 18.4 | 11.5 | W > H | Bacteroidetes | Flavobacteriia | Flavobacteriales | Flavobacteriaceae | Flavobacterium |
| Otu000114 | 14.6 | 1.32E-04 | 1.04E-02 | 1.1 | 0.0 | W > H | Bacteroidetes | Flavobacteriia | Flavobacteriales | Flavobacteriaceae | Flavobacterium |
| Otu000121 | 14.6 | 1.32E-04 | 1.04E-02 | 1.0 | 0.0 | W > H | Bacteroidetes | Flavobacteriia | Flavobacteriales | Flavobacteriaceae | Flavobacterium |
| Otu000256 | 14.6 | 1.33E-04 | 1.04E-02 | 1.9 | 0.0 | W > H | Planctomycetes | Planctomycetia | Planctomycetales | Planctomycetaceae | Planctomyces |
| Otu000204 | 14.6 | 1.33E-04 | 1.04E-02 | 2.9 | 0.0 | W > H | Proteobacteria | Betaproteobacteria | Burkholderiales | Comamonadaceae | unclassified |
| Otu000145 | 14.6 | 1.34E-04 | 1.04E-02 | 3.4 | 0.0 | W > H | Proteobacteria | Alphaproteobacteria | Sphingomonadales | Sphingomonadaceae | unclassified |
| Otu000011 | 14.3 | 1.57E-04 | 1.18E-02 | 153.5 | 1.6 | W > H | unclassified | unclassified | unclassified | unclassified | unclassified |
| Otu000016 | 13.6 | 2.21E-04 | 1.56E-02 | 6.1 | 0.0 | W > H | Proteobacteria | Gammaproteobacteria | Legionellales | Coxiellaceae | Rickettsiella |
| Otu000280 | 13.5 | 2.40E-04 | 1.62E-02 | 0.9 | 0.0 | W > H | Proteobacteria | Alphaproteobacteria | Sphingomonadales | Sphingomonadaceae | unclassified |
| Otu000219 | 13.5 | 2.41E-04 | 1.62E-02 | 1.5 | 0.0 | W > H | Proteobacteria | Alphaproteobacteria | Sphingomonadales | unclassified | unclassified |
| Otu000242 | 13.5 | 2.41E-04 | 1.62E-02 | 2.2 | 0.0 | W > H | Planctomycetes | Planctomycetia | Pirellulales | Pirellulaceae | A17 |
| Otu000228 | 13.5 | 2.42E-04 | 1.62E-02 | 3.8 | 0.0 | W > H | Bacteroidetes | unclassified | unclassified | unclassified | unclassified |
| Otu000253 | 13.5 | 2.42E-04 | 1.62E-02 | 4.3 | 0.0 | W > H | SR1 | unclassified | unclassified | unclassified | unclassified |
| Otu001476 | 12.5 | 4.12E-04 | 2.58E-02 | 0.7 | 0.0 | W > H | unclassified | unclassified | unclassified | unclassified | unclassified |
| Otu000233 | 12.4 | 4.30E-04 | 2.58E-02 | 1.3 | 0.0 | W > H | Proteobacteria | Alphaproteobacteria | Rhodobacterales | Rhodobacteraceae | unclassified |
| Otu000230 | 12.4 | 4.30E-04 | 2.58E-02 | 1.1 | 0.0 | W > H | Actinobacteria | Actinobacteria | Actinomycetales | Mycobacteriaceae | Mycobacterium |
| Otu000264 | 12.4 | 4.30E-04 | 2.58E-02 | 1.4 | 0.0 | W > H | Planctomycetes | Planctomycetia | Pirellulales | Pirellulaceae | unclassified |
| Otu000320 | 12.4 | 4.31E-04 | 2.58E-02 | 1.1 | 0.0 | W > H | Proteobacteria | Alphaproteobacteria | Rhodospirillales | Rhodospirillaceae | Reyranella |
| Otu000275 | 12.4 | 4.31E-04 | 2.58E-02 | 9.4 | 0.0 | W > H | Firmicutes | Bacilli | Bacillales | Bacillaceae | Bacillus |
| Otu000654 | 12.4 | 4.32E-04 | 2.58E-02 | 1.1 | 0.0 | W > H | Actinobacteria | Actinobacteria | Actinomycetales | unclassified | unclassified |
| Otu000447 | 12.4 | 4.32E-04 | 2.58E-02 | 1.5 | 0.0 | W > H | Proteobacteria | Alphaproteobacteria | Rhizobiales | unclassified | unclassified |
| Otu000051 | 12.4 | 4.32E-04 | 2.58E-02 | 1.8 | 0.0 | W > H | Bacteroidetes | Flavobacteriia | Flavobacteriales | Flavobacteriaceae | Flavobacterium |

| | | | | | | | | | | | |
|---|---|---|---|---|---|---|---|---|---|---|---|
| Otu000130 | 12.5 | 4.14E-04 | 2.58E-02 | 8.9 | 0.0 | W > H | Proteobacteria | Alphaproteobacteria | Rickettsiales | Anaplasmataceae | Neorickettsia |
| Otu000283 | 11.9 | 5.70E-04 | 3.03E-02 | 1.2 | 0.3 | W > H | Proteobacteria | Alphaproteobacteria | Rhodobacterales | Rhodobacteraceae | Rhodobacter |
| Otu000652 | 11.4 | 7.44E-04 | 3.71E-02 | 0.6 | 0.0 | W > H | Proteobacteria | Alphaproteobacteria | Rhizobiales | unclassified | unclassified |
| Otu000922 | 11.3 | 7.58E-04 | 3.71E-02 | 1.2 | 0.0 | W > H | unclassified | unclassified | unclassified | unclassified | unclassified |
| Otu000355 | 11.3 | 7.58E-04 | 3.71E-02 | 1.9 | 0.0 | W > H | Proteobacteria | Alphaproteobacteria | Rhodospirillales | Rhodospirillaceae | Reyranella |
| Otu000421 | 11.3 | 7.60E-04 | 3.71E-02 | 1.0 | 0.0 | W > H | Verrucomicrobia | unclassified | unclassified | unclassified | unclassified |
| Otu000318 | 11.3 | 7.60E-04 | 3.71E-02 | 1.0 | 0.0 | W > H | Planctomycetes | Planctomycetia | Planctomycetales | Planctomycetaceae | Planctomyces |
| Otu000847 | 11.3 | 7.60E-04 | 3.71E-02 | 1.1 | 0.0 | W > H | Firmicutes | Clostridia | Clostridiales | unclassified | unclassified |
| Otu000445 | 11.3 | 7.61E-04 | 3.71E-02 | 3.4 | 0.0 | W > H | Tenericutes | Mollicutes | RsaHF231 | unclassified | unclassified |
| Otu000455 | 11.3 | 7.62E-04 | 3.71E-02 | 1.9 | 0.0 | W > H | Bacteroidetes | [Saprospirae] | [Saprospirales] | Saprospiraceae | unclassified |
| Otu000680 | 11.3 | 7.63E-04 | 3.71E-02 | 1.5 | 0.0 | W > H | Spirochaetes | Spirochaetes | [Borreliales] | [Borreliaceae] | Spironema |
| Otu000188 | 11.3 | 7.63E-04 | 3.71E-02 | 12.3 | 0.0 | W > H | Proteobacteria | Gammaproteobacteria | Xanthomonadales | Xanthomonadaceae | unclassified |
| Otu000105 | 11.3 | 7.64E-04 | 3.71E-02 | 2.1 | 0.0 | W > H | Proteobacteria | Gammaproteobacteria | Alteromonadales | OM60 | unclassified |
| Otu000064 | 11.3 | 7.86E-04 | 3.79E-02 | 21.6 | 0.0 | W > H | Firmicutes | Bacilli | Bacillales | unclassified | unclassified |
| Otu000396 | 11.2 | 8.29E-04 | 3.97E-02 | 1.6 | 0.0 | W > H | Proteobacteria | Alphaproteobacteria | Rhodobacterales | Rhodobacteraceae | Rhodobacter |
| Otu000529 | 23.1 | 1.51E-06 | 3.37E-04 | 0.0 | 8.2 | H > W | Firmicutes | Bacilli | Lactobacillales | Leuconostocaceae | Leuconostoc |
| Otu000610 | 23.1 | 1.51E-06 | 3.37E-04 | 0.0 | 8.0 | H > W | Firmicutes | Bacilli | Lactobacillales | Lactobacillaceae | Lactobacillus |
| Otu000235 | 23.1 | 1.52E-06 | 3.37E-04 | 0.0 | 16.4 | H > W | Proteobacteria | Gammaproteobacteria | Vibrionales | Vibrionaceae | Photobacterium |
| Otu000565 | 23.1 | 1.52E-06 | 3.37E-04 | 0.0 | 6.5 | H > W | Firmicutes | Bacilli | Lactobacillales | Lactobacillaceae | Lactobacillus |
| Otu000049 | 22.9 | 1.71E-06 | 3.55E-04 | 0.1 | 175.6 | H > W | Firmicutes | Bacilli | Lactobacillales | Leuconostocaceae | Weissella |
| Otu000073 | 21.5 | 3.56E-06 | 6.50E-04 | 0.2 | 77.4 | H > W | Firmicutes | Bacilli | Lactobacillales | Streptococcaceae | Streptococcus |
| Otu000174 | 16.3 | 5.37E-05 | 5.38E-03 | 0.0 | 9.2 | H > W | Firmicutes | Bacilli | Lactobacillales | Lactobacillaceae | Lactobacillus |
| Otu002558 | 15.6 | 7.94E-05 | 6.79E-03 | 0.0 | 1.0 | H > W | Firmicutes | Bacilli | Lactobacillales | Lactobacillaceae | Lactobacillus |
| Otu001118 | 15.6 | 7.97E-05 | 6.79E-03 | 0.0 | 3.9 | H > W | Fusobacteria | Fusobacteriia | Fusobacteriales | Fusobacteriaceae | unclassified |
| Otu000431 | 15.6 | 7.97E-05 | 6.79E-03 | 0.0 | 14.7 | H > W | Firmicutes | Bacilli | Lactobacillales | Lactobacillaceae | Lactobacillus |
| Otu000707 | 14.8 | 1.17E-04 | 9.74E-03 | 0.0 | 7.3 | H > W | Firmicutes | Bacilli | Lactobacillales | Streptococcaceae | Streptococcus |
| Otu000015 | 14.4 | 1.49E-04 | 1.14E-02 | 0.2 | 355.2 | H > W | Firmicutes | Clostridia | Clostridiales | Ruminococcaceae | unclassified |
| Otu000002 | 14.4 | 1.51E-04 | 1.15E-02 | 185.5 | 826.3 | H > W | Tenericutes | Mollicutes | Mycoplasmatales | Mycoplasmataceae | Mycoplasma |
| Otu000699 | 14.1 | 1.74E-04 | 1.29E-02 | 0.1 | 5.0 | H > W | Firmicutes | Bacilli | Lactobacillales | Lactobacillaceae | Lactobacillus |
| Otu001997 | 13.8 | 2.03E-04 | 1.45E-02 | 0.0 | 1.8 | H > W | Firmicutes | Clostridia | Clostridiales | unclassified | unclassified |

| | | | | | | | | | | | |
|---|---|---|---|---|---|---|---|---|---|---|---|
| Otu001863 | 13.8 | 2.03E-04 | 1.45E-02 | 0.0 | 1.5 | **H > W** | Firmicutes | Clostridia | Clostridiales | Clostridiaceae | Clostridium |
| Otu000753 | 13.8 | 2.03E-04 | 1.45E-02 | 0.0 | 4.3 | **H > W** | Firmicutes | Bacilli | Lactobacillales | Lactobacillaceae | Pediococcus |
| Otu000004 | 12.7 | 3.72E-04 | 2.46E-02 | 0.0 | 1.6 | **H > W** | Proteobacteria | Gammaproteobacteria | Pseudomonadales | Moraxellaceae | Acinetobacter |
| Otu000703 | 12.1 | 5.06E-04 | 2.72E-02 | 0.0 | 2.0 | **H > W** | Firmicutes | Bacilli | Lactobacillales | Lactobacillaceae | Lactobacillus |
| Otu000628 | 12.1 | 5.06E-04 | 2.72E-02 | 0.0 | 1.9 | **H > W** | Firmicutes | Bacilli | Lactobacillales | Streptococcaceae | Lactococcus |
| Otu000673 | 12.1 | 5.06E-04 | 2.72E-02 | 0.0 | 7.3 | **H > W** | Firmicutes | Bacilli | Lactobacillales | Lactobacillaceae | Lactobacillus |
| Otu001153 | 12.1 | 5.06E-04 | 2.72E-02 | 0.0 | 2.9 | **H > W** | Firmicutes | Bacilli | Lactobacillales | Lactobacillaceae | Lactobacillus |
| Otu001521 | 12.1 | 5.06E-04 | 2.72E-02 | 0.0 | 2.1 | **H > W** | Firmicutes | Clostridia | Clostridiales | [Tissierellaceae] | Tepidimicrobium |
| Otu000788 | 12.1 | 5.06E-04 | 2.72E-02 | 0.0 | 4.1 | **H > W** | Proteobacteria | Gammaproteobacteria | Vibrionales | unclassified | unclassified |
| Otu001669 | 12.1 | 5.06E-04 | 2.72E-02 | 0.0 | 2.6 | **H > W** | Firmicutes | Clostridia | Clostridiales | Peptostreptococcaceae | Peptostreptococcus |
| Otu001874 | 12.1 | 5.06E-04 | 2.72E-02 | 0.0 | 1.9 | **H > W** | Firmicutes | Clostridia | Clostridiales | Lachnospiraceae | unclassified |
| Otu000525 | 12.1 | 5.07E-04 | 2.72E-02 | 0.0 | 11.5 | **H > W** | Fusobacteria | Fusobacteriia | Fusobacteriales | Fusobacteriaceae | Fusobacterium |
| Otu000614 | 12.1 | 5.07E-04 | 2.72E-02 | 0.0 | 9.2 | **H > W** | Fusobacteria | Fusobacteriia | Fusobacteriales | Fusobacteriaceae | unclassified |
| Otu001184 | 12.1 | 5.07E-04 | 2.72E-02 | 0.0 | 4.5 | **H > W** | Proteobacteria | Gammaproteobacteria | Vibrionales | Vibrionaceae | Photobacterium |
| Otu001352 | 12.1 | 5.07E-04 | 2.72E-02 | 0.0 | 2.6 | **H > W** | Fusobacteria | Fusobacteriia | Fusobacteriales | Fusobacteriaceae | Fusobacterium |

**Table S4.** Differentially abundant OTUs identified between wild and hatchery fish in the skin.

| OTU | Test-Statistic | P | FDR_P | Wild mean abundance | Hatchery mean abundance | Direction | Phylum | Class | Order | Family | Genus |
|---|---|---|---|---|---|---|---|---|---|---|---|
| Otu000066 | 49.56 | 1.92E-12 | 1.19E-08 | 7.42 | 0.00 | W > H | Proteobacteria | Alphaproteobacteria | Caulobacterales | Caulobacteraceae | Mycoplana |
| Otu000041 | 41.80 | 1.01E-10 | 3.14E-07 | 9.53 | 0.09 | W > H | Verrucomicrobia | Verrucomicrobiae | Verrucomicrobiales | Verrucomicrobiaceae | Luteolibacter |
| Otu000060 | 39.04 | 4.15E-10 | 8.60E-07 | 11.40 | 0.21 | W > H | Proteobacteria | Alphaproteobacteria | Caulobacterales | Caulobacteraceae | Mycoplana |
| Otu000016 | 38.04 | 6.94E-10 | 1.08E-06 | 54.40 | 0.15 | W > H | Proteobacteria | Gammaproteobacteria | Legionellales | Coxiellaceae | Rickettsiella |
| Otu000027 | 36.46 | 1.56E-09 | 1.94E-06 | 24.49 | 2.41 | W > H | Proteobacteria | Betaproteobacteria | Burkholderiales | Oxalobacteraceae | Massilia |
| Otu000168 | 35.99 | 1.98E-09 | 2.05E-06 | 3.47 | 0.15 | W > H | Proteobacteria | Betaproteobacteria | unclassified | unclassified | unclassified |
| Otu000063 | 34.43 | 4.41E-09 | 3.92E-06 | 9.89 | 0.76 | W > H | Proteobacteria | Alphaproteobacteria | Rhizobiales | Rhizobiaceae | Agrobacterium |
| Otu000072 | 32.64 | 1.11E-08 | 8.61E-06 | 6.18 | 0.24 | W > H | Proteobacteria | Alphaproteobacteria | Sphingomonadales | Sphingomonadaceae | Sphingomonas |
| Otu000065 | 32.23 | 1.37E-08 | 9.45E-06 | 5.27 | 0.53 | W > H | Proteobacteria | Alphaproteobacteria | Rhodobacterales | Rhodobacteraceae | Rhodobacter |
| Otu000169 | 31.96 | 1.57E-08 | 9.78E-06 | 3.96 | 0.00 | W > H | Actinobacteria | Actinobacteria | Actinomycetales | Micrococcaceae | Arthrobacter |
| Otu000020 | 30.85 | 2.79E-08 | 1.58E-05 | 4.89 | 0.59 | W > H | Proteobacteria | Gammaproteobacteria | Enterobacteriales | Enterobacteriaceae | unclassified |
| Otu000084 | 28.90 | 7.62E-08 | 3.95E-05 | 12.76 | 0.00 | W > H | Thermi | Deinococci | Deinococcales | Deinococcaceae | Deinococcus |
| Otu000076 | 27.46 | 1.60E-07 | 7.68E-05 | 10.82 | 0.00 | W > H | Proteobacteria | Betaproteobacteria | Burkholderiales | Oxalobacteraceae | unclassified |
| Otu000112 | 26.84 | 2.21E-07 | 9.80E-05 | 2.51 | 0.03 | W > H | Bacteroidetes | Cytophagia | Cytophagales | Cytophagaceae | Leadbetterella |
| Otu000161 | 26.56 | 2.56E-07 | 9.93E-05 | 3.73 | 0.15 | W > H | Bacteroidetes | Sphingobacteriia | Sphingobacteriales | Sphingobacteriaceae | Pedobacter |
| Otu000054 | 26.61 | 2.49E-07 | 9.93E-05 | 4.71 | 0.50 | W > H | Proteobacteria | Betaproteobacteria | Methylophilales | Methylophilaceae | Methylotenera |
| Otu000010 | 25.01 | 5.71E-07 | 2.09E-04 | 77.29 | 33.09 | W > H | Proteobacteria | Betaproteobacteria | Burkholderiales | Comamonadaceae | Rhodoferax |
| Otu000107 | 24.65 | 6.87E-07 | 2.25E-04 | 2.82 | 0.00 | W > H | Verrucomicrobia | Verrucomicrobiae | Verrucomicrobiales | Verrucomicrobiaceae | unclassified |
| Otu000179 | 24.65 | 6.87E-07 | 2.25E-04 | 3.71 | 0.00 | W > H | Proteobacteria | Alphaproteobacteria | Sphingomonadales | Sphingomonadaceae | Kaistobacter |
| Otu000118 | 24.39 | 7.88E-07 | 2.41E-04 | 4.93 | 0.12 | W > H | Proteobacteria | Gammaproteobacteria | Xanthomonadales | Xanthomonadaceae | Stenotrophomonas |
| Otu000013 | 24.32 | 8.15E-07 | 2.41E-04 | 69.36 | 25.44 | W > H | Proteobacteria | Betaproteobacteria | Burkholderiales | Comamonadaceae | unclassified |
| Otu000120 | 24.18 | 8.77E-07 | 2.48E-04 | 5.40 | 0.74 | W > H | Proteobacteria | Alphaproteobacteria | Caulobacterales | Caulobacteraceae | Brevundimonas |
| Otu000087 | 23.47 | 1.27E-06 | 3.44E-04 | 2.60 | 0.06 | W > H | Proteobacteria | Alphaproteobacteria | Rhizobiales | unclassified | unclassified |
| Otu000096 | 22.65 | 1.94E-06 | 5.03E-04 | 2.27 | 0.18 | W > H | Proteobacteria | Betaproteobacteria | Burkholderiales | Comamonadaceae | unclassified |
| Otu000026 | 21.83 | 2.98E-06 | 7.42E-04 | 13.67 | 2.59 | W > H | Proteobacteria | Alphaproteobacteria | Rhodobacterales | Rhodobacteraceae | Rhodobacter |
| Otu000068 | 20.69 | 5.41E-06 | 1.29E-03 | 4.73 | 0.00 | W > H | Cyanobacteria | Oscillatoriophycideae | Chroococcales | Xenococcaceae | Chroococcidiopsis |

| | | | | | | | | | | | |
|---|---|---|---|---|---|---|---|---|---|---|---|
| Otu000136 | 19.47 | 1.02E-05 | 2.22E-03 | 1.07 | 0.00 | W > H | Verrucomicrobia | Verrucomicrobiae | Verrucomicrobiales | Verrucomicrobiaceae | unclassified |
| Otu000147 | 19.45 | 1.03E-05 | 2.22E-03 | 1.76 | 0.00 | W > H | Verrucomicrobia | Verrucomicrobiae | Verrucomicrobiales | Verrucomicrobiaceae | unclassified |
| Otu000266 | 19.45 | 1.03E-05 | 2.22E-03 | 1.47 | 0.00 | W > H | Bacteroidetes | Flavobacteriia | Flavobacteriales | Weeksellaceae | Chryseobacterium |
| Otu000044 | 19.26 | 1.14E-05 | 2.37E-03 | 8.58 | 1.68 | W > H | Firmicutes | Bacilli | Bacillales | Exiguobacteraceae | Exiguobacterium |
| Otu000028 | 19.19 | 1.18E-05 | 2.37E-03 | 51.02 | 0.12 | W > H | Proteobacteria | unclassified | unclassified | unclassified | unclassified |
| Otu000091 | 18.26 | 1.93E-05 | 3.52E-03 | 10.87 | 0.62 | W > H | Bacteroidetes | Flavobacteriia | Flavobacteriales | Weeksellaceae | Chryseobacterium |
| Otu000114 | 17.94 | 2.28E-05 | 4.06E-03 | 3.38 | 0.47 | W > H | Bacteroidetes | Flavobacteriia | Flavobacteriales | Flavobacteriaceae | Flavobacterium |
| Otu000099 | 17.37 | 3.07E-05 | 5.30E-03 | 8.27 | 2.06 | W > H | Proteobacteria | Alphaproteobacteria | Rhizobiales | Methylobacteriaceae | Methylobacterium |
| Otu000426 | 17.03 | 3.68E-05 | 5.49E-03 | 1.53 | 0.00 | W > H | Bacteroidetes | Cytophagia | Cytophagales | Cytophagaceae | Hymenobacter |
| Otu000231 | 17.02 | 3.70E-05 | 5.49E-03 | 1.62 | 0.00 | W > H | Proteobacteria | Alphaproteobacteria | Rhizobiales | Hyphomicrobiaceae | Devosia |
| Otu000220 | 17.02 | 3.71E-05 | 5.49E-03 | 2.84 | 0.00 | W > H | Acidobacteria | Acidobacteria-6 | iii1-15 | unclassified | unclassified |
| Otu000145 | 17.07 | 3.61E-05 | 5.49E-03 | 1.33 | 0.03 | W > H | Proteobacteria | Alphaproteobacteria | Sphingomonadales | Sphingomonadaceae | unclassified |
| Otu000189 | 17.12 | 3.51E-05 | 5.49E-03 | 3.93 | 1.09 | W > H | Proteobacteria | Gammaproteobacteria | Pseudomonadales | Moraxellaceae | Enhydrobacter |
| Otu000184 | 16.44 | 5.02E-05 | 7.09E-03 | 2.62 | 0.44 | W > H | Proteobacteria | Alphaproteobacteria | Sphingomonadales | Sphingomonadaceae | Sphingomonas |
| Otu000140 | 16.31 | 5.39E-05 | 7.45E-03 | 2.62 | 0.29 | W > H | Firmicutes | Bacilli | Bacillales | unclassified | unclassified |
| Otu000135 | 15.87 | 6.79E-05 | 8.86E-03 | 1.53 | 0.00 | W > H | Proteobacteria | Alphaproteobacteria | Rhizobiales | unclassified | unclassified |
| Otu000105 | 15.87 | 6.80E-05 | 8.86E-03 | 1.09 | 0.00 | W > H | Proteobacteria | Gammaproteobacteria | Alteromonadales | OM60 | unclassified |
| Otu000146 | 15.85 | 6.84E-05 | 8.86E-03 | 3.58 | 0.00 | W > H | Bacteroidetes | Cytophagia | Cytophagales | Cytophagaceae | Hymenobacter |
| Otu000153 | 15.73 | 7.32E-05 | 9.29E-03 | 4.87 | 1.21 | W > H | Proteobacteria | Alphaproteobacteria | Rhizobiales | Methylobacteriaceae | Methylobacterium |
| Otu000043 | 14.83 | 1.17E-04 | 1.33E-02 | 6.49 | 0.09 | W > H | Proteobacteria | Gammaproteobacteria | Alteromonadales | OM60 | unclassified |
| Otu000348 | 14.73 | 1.24E-04 | 1.36E-02 | 1.67 | 0.00 | W > H | Firmicutes | Bacilli | Bacillales | Planococcaceae | Paenisporosarcina |
| Otu000148 | 14.72 | 1.25E-04 | 1.36E-02 | 1.51 | 0.00 | W > H | Verrucomicrobia | Verrucomicrobiae | Verrucomicrobiales | Verrucomicrobiaceae | Luteolibacter |
| Otu000165 | 13.89 | 1.94E-04 | 2.01E-02 | 5.24 | 0.47 | W > H | Proteobacteria | Gammaproteobacteria | Oceanospirillales | Halomonadaceae | Halomonas |
| Otu000046 | 13.67 | 2.17E-04 | 2.22E-02 | 9.62 | 3.35 | W > H | Proteobacteria | Betaproteobacteria | Burkholderiales | Comamonadaceae | Hydrogenophaga |
| Otu000131 | 13.62 | 2.24E-04 | 2.22E-02 | 1.58 | 0.00 | W > H | Proteobacteria | Betaproteobacteria | SC-I-84 | unclassified | unclassified |
| Otu000201 | 13.61 | 2.25E-04 | 2.22E-02 | 2.47 | 0.00 | W > H | Proteobacteria | Alphaproteobacteria | Sphingomonadales | Sphingomonadaceae | Sphingopyxis |
| Otu000167 | 13.50 | 2.38E-04 | 2.32E-02 | 1.78 | 0.03 | W > H | Planctomycetes | Planctomycetia | Pirellulales | Pirellulaceae | unclassified |
| Otu000233 | 13.38 | 2.54E-04 | 2.39E-02 | 1.22 | 0.03 | W > H | Proteobacteria | Alphaproteobacteria | Rhodobacterales | Rhodobacteraceae | unclassified |
| Otu000083 | 13.41 | 2.51E-04 | 2.39E-02 | 11.04 | 1.18 | W > H | Actinobacteria | Actinobacteria | Actinomycetales | Nocardiaceae | Rhodococcus |
| Otu000182 | 13.12 | 2.92E-04 | 2.49E-02 | 3.02 | 0.76 | W > H | Firmicutes | Bacilli | Bacillales | Planococcaceae | unclassified |

| | | | | | | | | | | | |
|---|---|---|---|---|---|---|---|---|---|---|---|
| Otu000324 | 13.04 | 3.05E-04 | 2.57E-02 | 0.98 | 0.09 | W > H | Bacteroidetes | Cytophagia | Cytophagales | Cytophagaceae | Emticicia |
| Otu000635 | 12.56 | 3.95E-04 | 2.79E-02 | 0.78 | 0.00 | W > H | Bacteroidetes | Cytophagia | Cytophagales | Cytophagaceae | Leadbetterella |
| Otu000256 | 12.55 | 3.96E-04 | 2.79E-02 | 0.93 | 0.00 | W > H | Planctomycetes | Planctomycetia | Planctomycetales | Planctomycetaceae | Planctomyces |
| Otu000270 | 12.55 | 3.97E-04 | 2.79E-02 | 0.96 | 0.00 | W > H | Actinobacteria | Acidimicrobiia | Acidimicrobiales | C111 | unclassified |
| Otu000280 | 12.55 | 3.97E-04 | 2.79E-02 | 0.91 | 0.00 | W > H | Proteobacteria | Alphaproteobacteria | Sphingomonadales | Sphingomonadaceae | unclassified |
| Otu000180 | 12.54 | 3.98E-04 | 2.79E-02 | 1.07 | 0.00 | W > H | Proteobacteria | Alphaproteobacteria | Sphingomonadales | Sphingomonadaceae | unclassified |
| Otu000612 | 12.54 | 3.98E-04 | 2.79E-02 | 1.22 | 0.00 | W > H | Firmicutes | Bacilli | Bacillales | Planococcaceae | Paenisporosarcina |
| Otu000693 | 12.54 | 3.98E-04 | 2.79E-02 | 1.02 | 0.00 | W > H | Actinobacteria | Actinobacteria | Actinomycetales | Micrococcaceae | unclassified |
| Otu000185 | 12.54 | 3.98E-04 | 2.79E-02 | 1.11 | 0.00 | W > H | Proteobacteria | Gammaproteobacteria | Thiotrichales | Piscirickettsiaceae | unclassified |
| Otu000134 | 12.54 | 3.99E-04 | 2.79E-02 | 1.98 | 0.00 | W > H | Verrucomicrobia | Spartobacteria | Chthoniobacterales | Chthoniobacteraceae | Chthoniobacter |
| Otu000216 | 12.54 | 3.99E-04 | 2.79E-02 | 5.56 | 0.00 | W > H | Proteobacteria | Betaproteobacteria | Neisseriales | Neisseriaceae | Vogesella |
| Otu000518 | 12.54 | 3.99E-04 | 2.79E-02 | 1.73 | 0.00 | W > H | TM7 | TM7-3 | EW055 | unclassified | unclassified |
| Otu000098 | 12.54 | 3.99E-04 | 2.79E-02 | 23.33 | 0.03 | W > H | Proteobacteria | Betaproteobacteria | Burkholderiales | Alcaligenaceae | unclassified |
| Otu000186 | 12.74 | 3.57E-04 | 2.79E-02 | 2.38 | 0.12 | W > H | Planctomycetes | Planctomycetia | Pirellulales | Pirellulaceae | unclassified |
| Otu000055 | 12.58 | 3.90E-04 | 2.79E-02 | 36.49 | 3.41 | W > H | Firmicutes | Bacilli | Bacillales | Staphylococcaceae | Staphylococcus |
| Otu000097 | 12.63 | 3.79E-04 | 2.79E-02 | 9.73 | 2.24 | W > H | Actinobacteria | Actinobacteria | Actinomycetales | Micrococcaceae | Micrococcus |
| Otu000240 | 12.44 | 4.20E-04 | 2.90E-02 | 1.60 | 0.03 | W > H | Bacteroidetes | Cytophagia | Cytophagales | Cytophagaceae | Dyadobacter |
| Otu000451 | 12.27 | 4.61E-04 | 3.15E-02 | 1.38 | 0.03 | W > H | Actinobacteria | Actinobacteria | Actinomycetales | Intrasporangiaceae | Phycicoccus |
| Otu000077 | 12.23 | 4.71E-04 | 3.19E-02 | 5.73 | 0.35 | W > H | Proteobacteria | Betaproteobacteria | Procabacteriales | Procabacteriaceae | unclassified |
| Otu000133 | 11.83 | 5.82E-04 | 3.81E-02 | 5.60 | 0.62 | W > H | Actinobacteria | Actinobacteria | Actinomycetales | Nocardiaceae | Rhodococcus |
| Otu000191 | 11.73 | 6.15E-04 | 3.98E-02 | 1.42 | 0.12 | W > H | Firmicutes | Bacilli | Bacillales | Bacillaceae | Bacillus |
| Otu000075 | 11.63 | 6.48E-04 | 4.03E-02 | 2.62 | 0.47 | W > H | Proteobacteria | Betaproteobacteria | Burkholderiales | Comamonadaceae | Limnohabitans |
| Otu000009 | 11.64 | 6.46E-04 | 4.03E-02 | 65.13 | 11.88 | W > H | Proteobacteria | Gammaproteobacteria | Pseudomonadales | Moraxellaceae | Acinetobacter |
| Otu000414 | 11.50 | 6.96E-04 | 4.07E-02 | 1.24 | 0.00 | W > H | Bacteroidetes | Cytophagia | Cytophagales | Cytophagaceae | Hymenobacter |
| Otu000113 | 11.50 | 6.97E-04 | 4.07E-02 | 0.87 | 0.00 | W > H | Proteobacteria | Gammaproteobacteria | Xanthomonadales | Sinobacteraceae | unclassified |
| Otu000467 | 11.50 | 6.97E-04 | 4.07E-02 | 1.78 | 0.00 | W > H | Actinobacteria | Actinobacteria | Actinomycetales | Nocardioidaceae | Nocardioides |
| Otu000719 | 11.49 | 6.99E-04 | 4.07E-02 | 0.93 | 0.00 | W > H | Cyanobacteria | Synechococcophycideae | Pseudanabaenales | Pseudanabaenaceae | unclassified |
| Otu000128 | 11.49 | 7.00E-04 | 4.07E-02 | 1.56 | 0.00 | W > H | Proteobacteria | unclassified | unclassified | unclassified | unclassified |
| Otu000286 | 11.49 | 7.00E-04 | 4.07E-02 | 1.02 | 0.00 | W > H | Acidobacteria | Acidobacteria-6 | iii1-15 | unclassified | unclassified |
| Otu000274 | 11.49 | 7.01E-04 | 4.07E-02 | 2.16 | 0.00 | W > H | Proteobacteria | Alphaproteobacteria | Rhodospirillales | Acetobacteraceae | Roseomonas |

| | | | | | | | | | | | |
|---|---|---|---|---|---|---|---|---|---|---|---|
| Otu000061 | 11.43 | 7.21E-04 | 4.15E-02 | 10.78 | 2.82 | W > H | Firmicutes | Bacilli | Lactobacillales | Streptococcaceae | Streptococcus |
| Otu000111 | 11.27 | 7.88E-04 | 4.50E-02 | 4.67 | 0.65 | W > H | Proteobacteria | Alphaproteobacteria | Rhizobiales | Bradyrhizobiaceae | Bradyrhizobium |
| Otu000048 | 11.25 | 7.98E-04 | 4.51E-02 | 3.36 | 0.18 | W > H | Proteobacteria | Betaproteobacteria | Neisseriales | Neisseriaceae | Deefgea |
| Otu000004 | 18.99 | 1.31E-05 | 2.55E-03 | 3.11 | 116.26 | H> W | Proteobacteria | Gammaproteobacteria | Pseudomonadales | Moraxellaceae | Acinetobacter |
| Otu000395 | 18.33 | 1.86E-05 | 3.50E-03 | 0.00 | 2.44 | H> W | Proteobacteria | Gammaproteobacteria | Legionellales | Legionellaceae | Legionella |
| Otu000030 | 17.21 | 3.35E-05 | 5.49E-03 | 0.24 | 38.68 | H> W | Proteobacteria | Betaproteobacteria | unclassified | unclassified | unclassified |
| Otu000548 | 16.58 | 4.66E-05 | 6.73E-03 | 0.00 | 1.76 | H> W | Proteobacteria | Gammaproteobacteria | unclassified | unclassified | unclassified |
| Otu000770 | 14.88 | 1.14E-04 | 1.32E-02 | 0.00 | 0.59 | H> W | Proteobacteria | Alphaproteobacteria | Sphingomonadales | Sphingomonadaceae | unclassified |
| Otu001125 | 14.88 | 1.15E-04 | 1.32E-02 | 0.00 | 0.71 | H> W | Proteobacteria | Gammaproteobacteria | unclassified | unclassified | unclassified |
| Otu000217 | 14.88 | 1.15E-04 | 1.32E-02 | 0.00 | 2.76 | H> W | Bacteroidetes | Cytophagia | Cytophagales | unclassified | unclassified |
| Otu000521 | 14.87 | 1.15E-04 | 1.32E-02 | 0.00 | 1.41 | H> W | TM7 | TM7-1 | unclassified | unclassified | unclassified |
| Otu000406 | 14.87 | 1.15E-04 | 1.32E-02 | 0.00 | 2.47 | H> W | unclassified | unclassified | unclassified | unclassified | unclassified |
| Otu000342 | 14.00 | 1.83E-04 | 1.93E-02 | 0.02 | 2.97 | H> W | TM7 | SC3 | unclassified | unclassified | unclassified |
| Otu000197 | 14.00 | 1.83E-04 | 1.93E-02 | 0.02 | 5.85 | H> W | Bacteroidetes | Cytophagia | Cytophagales | Cytophagaceae | Rudanella |
| Otu000664 | 13.22 | 2.78E-04 | 2.40E-02 | 0.00 | 0.76 | H> W | Bacteroidetes | Cytophagia | Cytophagales | Cytophagaceae | Leadbetterella |
| Otu000927 | 13.21 | 2.78E-04 | 2.40E-02 | 0.00 | 0.88 | H> W | Proteobacteria | unclassified | unclassified | unclassified | unclassified |
| Otu000980 | 13.21 | 2.78E-04 | 2.40E-02 | 0.00 | 0.71 | H> W | unclassified | unclassified | unclassified | unclassified | unclassified |
| Otu000978 | 13.21 | 2.78E-04 | 2.40E-02 | 0.00 | 0.76 | H> W | TM6 | SBRH58 | unclassified | unclassified | unclassified |
| Otu000863 | 13.21 | 2.78E-04 | 2.40E-02 | 0.00 | 0.91 | H> W | Chloroflexi | Anaerolineae | Caldilineales | Caldilineaceae | unclassified |
| Otu000806 | 13.21 | 2.78E-04 | 2.40E-02 | 0.00 | 1.06 | H> W | Proteobacteria | unclassified | unclassified | unclassified | unclassified |
| Otu000432 | 12.16 | 4.87E-04 | 3.26E-02 | 0.02 | 1.97 | H> W | Bacteroidetes | Cytophagia | Cytophagales | Cytophagaceae | Runella |
| Otu000012 | 12.11 | 5.02E-04 | 3.32E-02 | 0.80 | 9.35 | H> W | Proteobacteria | Gammaproteobacteria | Pseudomonadales | Pseudomonadaceae | Pseudomonas |
| Otu000436 | 11.64 | 6.44E-04 | 4.03E-02 | 0.07 | 1.88 | H> W | Bacteroidetes | Flavobacteriia | Flavobacteriales | Flavobacteriaceae | Flavobacterium |
| Otu000371 | 11.64 | 6.44E-04 | 4.03E-02 | 0.07 | 2.38 | H> W | TM7 | SC3 | unclassified | unclassified | unclassified |
